# DGAT1 is a lipid metabolism oncoprotein that enables cancer cells to accumulate fatty acid while avoiding lipotoxicity

**DOI:** 10.1101/2020.06.23.166603

**Authors:** Daniel J. Wilcock, Andrew P. Badrock, Rhys Owen, Melissa Guerin, Andrew D. Southam, Hannah Johnston, Samuel Ogden, Paul Fullwood, Joanne Watson, Harriet Ferguson, Jennifer Ferguson, Daniel A. Richardson, Gavin R. Lloyd, Andris Jankevics, Warwick B. Dunn, Claudia Wellbrock, Paul Lorigan, Craig Ceol, Chiara Francavilla, Michael P. Smith, Adam F. L. Hurlstone

**Author notes:** These authors contributed equally (Daniel J. Wilcock, Andrew P. Badrock). Co-senior authors (Andrew P. Badrock, Chiara Francavilla). To whom correspondence should be addressed: Dr Michael P. Smith, Michael Smith Building, The University of Manchester, Dover Street, Manchester M13 9PT, UK;: Dr Adam Hurlstone, Michael Smith Building, The University of Manchester, Dover Street, Manchester M13 9PT, UK.

## Abstract

Dysregulated cellular metabolism is a hallmark of cancer. As yet, few druggable oncoproteins directly responsible for this hallmark have been identified. Increased fatty acid acquisition allows cancer cells to meet their membrane biogenesis, ATP, and signaling needs. Excess fatty acids suppress growth factor signaling and cause oxidative stress in non-transformed cells, but surprisingly not in cancer cells. Molecules underlying this cancer adaptation may provide new drug targets. Here, we identify Diacylglycerol O-acyltransferase 1 (DGAT1), an enzyme integral to triacylglyceride synthesis and lipid droplet formation, as a frequently up-regulated oncoprotein allowing cancer cells to tolerate excess fatty acids. DGAT1 over-expression alone induced melanoma in zebrafish melanocytes, and co-operated with oncogenic BRAF or NRAS for more rapid melanoma formation. Mechanistically, DGAT1 stimulated melanoma cell growth through sustaining mTOR kinase–S6 kinase signaling and suppressed cell death by tempering fatty acid oxidation, thereby preventing accumulation of reactive oxygen species including lipid peroxides.

**SIGNIFICANCE:** We show that DGAT1 is a *bona fide* oncoprotein capable of inducing melanoma formation and co-operating with other known drivers of melanoma. DGAT1 facilitates enhanced fatty acid acquisition by melanoma cells through suppressing lipototoxicity. DGAT1 is also critical for maintaining S6K activity required for melanoma cell growth.

## INTRODUCTION

The continuous growth and survival of cancer cells is underpinned by dysregulated cellular metabolism (1). Dysregulated cellular metabolism further promotes cancer progression by reprogramming stromal cells, mediating evasion of immune responses, and promoting metastasis (2-4). A central facet of cancer cell metabolism is a shift in glucose usage away from oxidative phosphorylation towards biosynthetic reactions. Without compensation by fatty acid oxidation (FAO) and glutaminolysis, this shift would result in deficiencies in citric acid cycle intermediates needed for ATP production (1).

Although catabolism of fatty acids (FA) is needed to maintain ATP production, increased lipogenesis (lipid generation) is simultaneously required for cell membrane synthesis for which FA are principal building blocks (5). Furthermore, lipid signaling molecules such as phosphatidyl inositides, diacylglycerides (DAG), lysophosphatidic acid, and prostaglandins implicated in multiple cancer hallmarks (6) also require FA. To satisfy these competing demands for FA, cancer cells increase its supply. This can be achieved by initiation of *de novo* FA synthesis, normally a function of adipocytes and hepatocytes, but an acquired characteristic of many cancer cells driven by elevated FA synthase (FASN) expression (5). Alternatively, the additional FA needed can be scavenged from the blood circulation as well as from abutting adipose tissue. Cancer cells utilize secreted lipases such as lipoprotein lipase (LPL) to hydrolyze FA from circulating triglycerides and FA transporter proteins (FATP) such as CD36 and the SLC27 family of FATP and FA binding proteins (FABP) to facilitate uptake (7). Enhanced uptake of extracellular FA by cancer cells has been established to promote tumor growth and dissemination (8-10). Intracellular lipases can also be called on to liberate FA from intracellular lipid stores, both through lipophagy or otherwise (11). One such lipase, monoacylglycerol lipase (MAGL) is up-regulated in some aggressive cancers, wherein its suppression impedes tumor growth and metastasis (12).

Regardless of the mechanism of acquisition, high levels of free FA come at a cost to most cells. Non-adipose, untransformed cells that become overloaded with FA display a constellation of effects termed lipotoxicity, characterized by reduced insulin signaling (insulin resistance) and increased cell death (13). Multiple processes underlie lipotoxicity, including increased ceramide synthesis, dysregulation of phospholipid production that compromises mitochondrial and endoplasmic reticulum (ER) membrane integrity, induction of reactive oxygen species (ROS), and impairment of ATP generation (13). How, therefore, dysregulated metabolism favoring FA accumulation is tolerated by cancer cells, and in particular how the toxic by-products of rampant FA metabolism (principally reactive oxygen species [ROS] including lipid peroxides) are suppressed or neutralized, is currently unclear. Targeting the ability of cancer cells to manage potentially cytotoxic metabolites that arise from the rewiring of metabolic pathways is an intriguing therapeutic avenue warranting further exploration (1).

*Bona fide* oncoproteins (defined as gene products whose up-regulation or coding alteration contributes directly to neoplasia), for example BCR-ABL, BRAF^V600E^ or mutant EGFR, are highly desirable molecular targets for precision treatment in cancers. Very few druggable oncoproteins have been identified that are directly responsible for dysregulating cellular metabolism. Among metabolic enzymes, only point mutations in the isocitrate dehydrogenases IDH1 and IDH2 have been confirmed as oncogenic drivers, but these mutations are detected in only 0.9%-3% of all cancers (14) (Supplementary Figure 1A). While the gene encoding FASN is frequently amplified in cancer (Supplementary Figure 1A), FASN inhibitors have so far failed to gain clinical approval (15). Much of the metabolic reprogramming of cancer cells is achieved through up-regulation of transcription factors including HIF1, MYC, peroxisome proliferator-activated receptors (PPARs) and sterol regulatory element binding factors (SREBFs) (16,17). However, these either lack druggable pockets or have diverse physiological effects, detracting from their therapeutic potential. Thus, there is currently a dearth of metabolism oncoproteins to pursue as therapeutic targets.

The small, fresh-water, tropical fish *Danio rerio*, commonly known as zebrafish, has been embraced as a model for melanoma oncogene discovery and validation (18). Previously, using zebrafish melanoma models, we identified a progression-associated transcript signature enriched for lipid metabolism genes and we established enhanced FA uptake as a characteristic of melanoma, driven in part by LPL (19). Here, we identify frequent amplification and up-regulation of the gene encoding Diacylglycerol O-acyltransferase 1 (*DGAT1*) in melanoma, among other cancers. DGAT1 is an ER-resident enzyme that catalyzes the final step in triacylglyceride (TAG) synthesis from diacylglyceride (DAG) and FA. DGAT1 is required for the formation of lipid droplets (LD), cytosolic organelles comprising a core of neutral lipids (mainly triglycerides and sterol esters) delimited by a monolayer of phospholipids, found in multiple cell types (20). Furthermore, we demonstrate the ability of DGAT1 to induce melanoma formation in zebrafish, and that DGAT1 exerts its oncogenic effect through shielding melanoma cells from lipotoxicity, while enhancing cell growth through sustaining mTOR–S6K signaling. As DGAT1 has been mooted as a clinical target for combating obesity, several potent and selective small molecule inhibitors are already available for repurposing (21) and we demonstrate their ability to suppress melanoma cell growth and survival.

## RESULTS

### Amplification and up-regulation of *DGAT1* associates with poor prognosis in melanoma and multiple other cancers

CD36, FATP1 and LPL that increase FA uptake into melanoma cells (9,19,22) are rarely activated by point mutation, nor are the genes encoding them often amplified in cancer, albeit that *CD36* amplification is enriched in established cancer cell lines (Supplementary Figure 1A). Given the lack of metabolism oncoproteins to pursue as therapeutic targets and the significance of lipid metabolism for melanoma development (23), we further interrogated the lipid metabolism genes that we previously identified accompany progression of mutant RAS driven melanoma in zebrafish (19). We plotted their expression fold-change against association of expression of the human homologue with patient survival. *dgat1a*/*DGAT1* emerged as an outlier, being both highly up-regulated in zebrafish melanoma and significantly associated with reduced patient survival (Figure 1A, B). In contrast, expression of the gene encoding the functionally related, although structurally distinct, Dgat2/DGAT2 (20) did not change significantly nor associate with patient survival (Figure 1A, B), indicating a possible unique function for DGAT1 in melanoma. Furthermore, we confirmed elevated expression of human *DGAT1* in melanoma tumors relative to both skin and nevi (Figure 1C) and elevated DGAT1 protein in human melanoma cell lines relative to primary melanocytes irrespective of NRAS or BRAF mutational status (Figure 1D).

**Figure 1.**
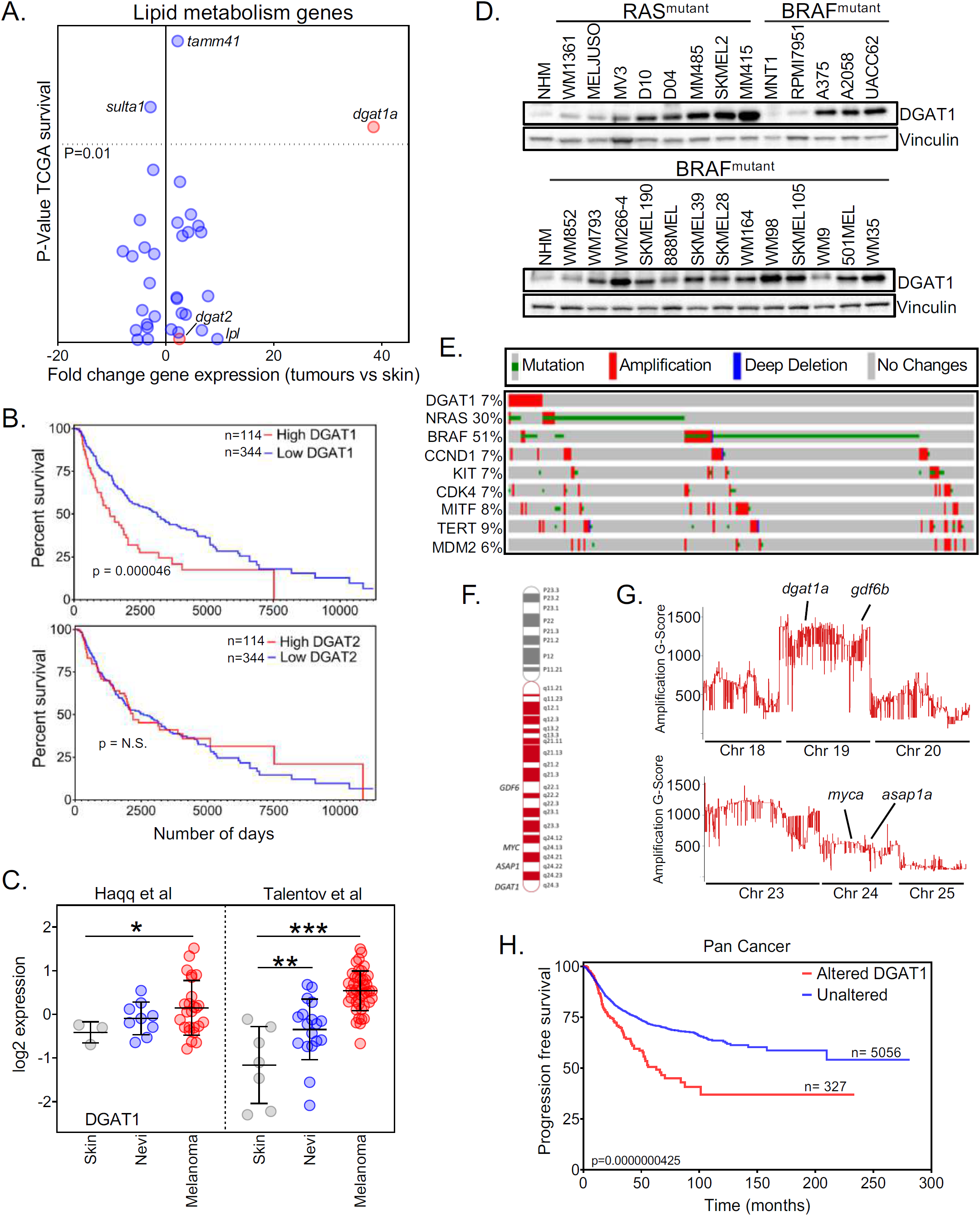
*DGAT1* amplification and up-regulation is associated with poor prognosis in melanoma. (a) Patient survival from TCGA melanoma cohort (25% top vs 75% bottom by mRNA abundance, Y-axis) *versus* fold-change in mRNA expression of lipid metabolism genes in zebrafish tumors (X-axis). (b) Kaplan-Meier survival plot comparing melanoma patients based on expression of *DGAT1* or *DGAT2* (top 25% *vs* bottom 75%, TCGA data set). (c) *DGAT1* relative gene expression in skin, nevi and melanoma tumors from indicated studies (Mean ± SD, n>3). (d) Protein expression of DGAT1 and Vinculin (loading control) (NHM-Normal Human Melanocytes). (e) Genetic alterations in the TCGA firehose legacy melanoma data set (counting only samples with CNV data) obtained from cBioPortal. (f) Schematic depicting human chromosome 8, the amplified arm (red), and known/putative melanoma oncogenes within this region. (g) G-Score of amplified regions of zebrafish chromosomes found in *BRAF*^*V600E*^-positive; *tp53* mutant tumors indicating the position of presumed melanoma oncogene homologues. (h) Kaplan-Meier progression free survival plot comparing patients across multiple cancer types based on *DGAT1* amplification.

We next considered what might underlie DGAT1 up-regulation in melanoma. Visualization of structural alterations of the *DGAT1* gene in The Cancer Genome Atlas firehose legacy cutaneous melanoma data set using cBioPortal revealed significant focal amplification (as defined by the stringent GISTIC 2.0 algorithm) in up to 7 % of melanoma cases with available copy number variation (CNV) data (but revealed no microscale-aberrations). This frequency of focal amplification was comparable to that for other recognized melanoma oncogenes (namely *CCND1, KIT, CDK4, MITF, TERT* and *MDM2*; Figure 1E). An extra copy of the long-arm of chromosome 8 (8q), which contains the *DGAT1* locus, but also other putative melanoma oncogenes *ASAP1*/*DDEF, MYC* and *GDF6* (24-26) (Figure 1F), has been observed in approximately 30 % of melanoma cases (27). Consistently, *DGAT1, ASAP1, MYC* and *GDF6* are co-amplified in melanoma and other cancers (Supplementary Figure 1B, C), although there were examples where each alone is amplified. However, of these four, high *DGAT1* mRNA expression displayed the strongest association with reduced patient survival (Figure 1B and Supplementary Figure 1D).

Further arguing for the significance of *DGAT1* amplification in melanoma development, our previous comparative oncogenomic analysis (24) uncovered amplification of *dgat1a* together with *gdf6b* (both on chromosome 19) in oncogenic-BRAF driven zebrafish melanoma, concomitant with up-regulation of *dgat1a* and *gdf6b* mRNA (Figure 1G and Supplementary Table 1). In contrast, neither *dgat1b* nor *gdf6a* on chromosome 16, nor any *myc* or *asap1* paralogues (on chromosomes 2 and 24) were amplified or up-regulated (Figure 1G, Supplementary Figure 1E, and Supplementary Table 1).

Finally, we found *DGAT1* to be amplified in other human cancers arising from distinct cell lineages, a feature of other well-characterized oncogenes, most notably in up to 26% of cases of ovarian cancer (Supplementary Figure 1F). Strikingly, *DGAT1* amplification was associated with significantly poorer progression-free survival across multiple cancer types (Figure 1H). Thus, from a cancer genomics perspective, *DGAT1* exhibits several hallmarks of an oncogene.

### Dgat1 functions as an oncoprotein in zebrafish melanocytes

In human melanoma, *DGAT1* amplification was observed to co-occur with *BRAF* and *NRAS* modification, but also independently of either (Figure 2A). To elucidate further the oncogenic potential of *DGAT1*, we utilized a melanocyte rescue and lineage restricted expression system as described previously (24,28). Remarkably, we found that Dgat1a over-expression in zebrafish melanocytes lacking functional p53 was sufficient to induce melanoma (Figure 2B), an outcome that we have only previously observed using the potent well-established oncogenes RAS and BRAF. Moreover, Dgat1a over-expression cooperated with both oncogenic BRAF and NRAS to accelerate the development of nodular tumors (Figure 2C-E and Supplementary Figure 2A). Thus, in zebrafish, Dgat1 behaves as a melanoma oncoprotein. Significantly, Dgat2 over-expression was indistinguishable from the non-oncogenic EGFP control (Figure 2D), indicating that Dgat1-mediated tumorigenesis is specific and cannot be replicated by the functionally related Dgat2.

**Figure 2.**
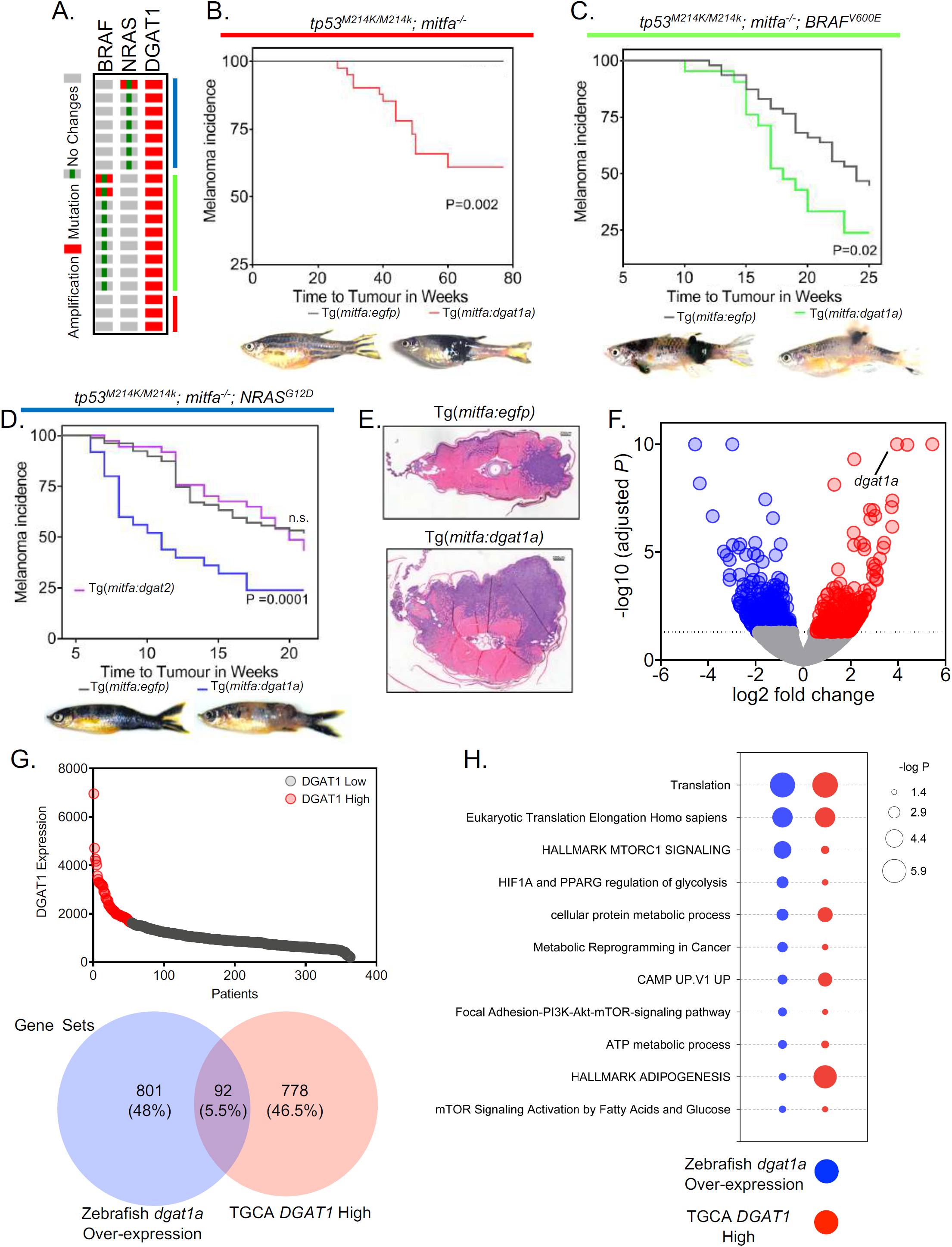
Dgat1 functions as an oncoprotein in zebrafish. (a) *DGAT1* amplification distribution in melanoma. (b) Kaplan-Meier plot of melanoma tumor nodule incidence in EGFP control or Dgat1a over-expressing animals on the *tp53* mutant; *nacre* genetic background. Representative images shown for EGFP and Dgat1a positive animals at 54 and 76 weeks post-fertilisation respectively. (c) as for (b) but on the transgenic *mitfa:BRAF*^*V600E*^; *tp53* mutant; *nacre* genetic background. Representative images are shown at 12 weeks post fertilisation. (d) as for (c) but on the transgenic *mitfa:NRAS*^*G12D*^; *tp53* mutant; *nacre* genetic background, also shown the effect of Dgat2 over-expression. (e) Hematoxylin and eosin stained transverse sections of EGFP expressing or Dgat1a over-expressing melanoma on the transgenic *mitfa*:*NRAS*^*G12D*^; *tp53* mutant; *nacre* genetic background. (f) Volcano plot of all genes from RNA-seq data from NRAS^G12D^-positive EGFP-expressing (n=3) and from NRAS^G12D^-positive Dgat1a over-expressing (n=5) tumors. Fold-change calculated comparing Dgat1a over-expressing tumors to EGFP-expressing control tumors. (g) *DGAT1* expression in TGCA pan cancer atlas RNA-seq data set filtered for NRAS^mut^ melanoma patients (upper). Venn diagram of significantly enriched gene sets after GSEA of significantly regulated genes comparing either NRAS^G12D^-positive EGFP-expressing (n=3) and NRAS^G12D^-positive Dgat1a over-expressing (n=5) tumors or comparing DGAT1 mRNA high tumors (12/72) to the rest of the NRAS^mut^ TGCA dataset (60/72). (h) Selection of 11 pathways from the Venn diagram intersect is shown in the dot-plot. Dot size represents the *P* value.

To enable the development of hypotheses explaining how *DGAT1* is a potent oncogene, we investigated the gene expression profile of melanoma tumors over-expressing Dgat1a, utilizing RNA sequencing (RNA-seq). RNA-seq revealed an average 5.5-fold increase in *dgat1a* mRNA expression compared to control EGFP-expressing tumors (Figure 2F). Principal component analysis (Supplementary Figure 2B) and. hierarchical clustering (Supplementary Figure 2C) revealed patterns of gene expression clearly distinguishing Dgat1a over-expressing tumors from EGFP-expressing tumors. To pinpoint the biological processes providing context for selection of DGAT1 up-regulation or that are potentially driven by DGAT1 up-regulation, we compared the gene sets enriched in Dgat1a over-expressing tumors to those enriched in the TCGA NRAS^mut^ DGAT1^high^ melanoma cohort compared to the rest of the NRAS^mut^ tumor samples in the dataset (Figure 2G and Supplementary Figure 2E and Supplementary Table 2). The overlap in enriched gene sets was only modest (Figure 2G) consistent with very few differentially expressed genes being shared in common (Supplementary Figure 2F). However, this was not altogether unexpected given that the genes whose expression most strongly correlated with *DGAT1* expression in human tumors consistently mapped to chromosome 8q (data not shown) and were therefore likely co-amplified with DGAT1, while *dgat1a* up-regulation in the zebrafish was driven by randomly integrated transgene combined with expression of endogenous *dgat1a* on a linkage group distinct from human 8q. Nonetheless, we hypothesized that the conserved gene sets (Supplementary Table 2) would be the most relevant to understanding the oncogenic role of *DGAT1*. Intriguingly, numerous of these conserved gene sets indicated enhanced activation of mTOR signaling and protein translation, as well as enhanced glycolysis and lipogenesis (Figure 2H), collectively indicative of increased insulin sensitivity in tumors with up-regulated *dgat1a*/*DGAT1*. From this analysis, therefore, we hypothesize that DGAT1 up-regulation modulates several biochemical hallmarks of cancer which are conserved in both zebrafish and human melanoma.

### DGAT1 enzymatic activity facilitates ribosomal protein S6-kinase (S6K)-stimulated growth

To ascertain which of the key oncogenic pathways highlighted by RNA-seq analysis of tumors may be directly dependent on DGAT1, we examined the initial signaling changes that occur when DGAT1 activity is suppressed, by performing unbiased quantitative mass-spectrometry (MS)-based phospho-proteomics. We analysed phosphorylated peptides from Stable Isotope Labelling in Cell Culture (SILAC)-labelled DGAT1^high^ A375 melanoma cells following 4h treatment with A922500. This revealed that DGAT1 activity affects cell growth and division signaling at mTOR, S6K and CDK1 signaling nodes (Figure 3A and Supplementary Figure 3A), mirroring the enrichment of the mTOR and protein translation signatures revealed by gene set enrichment analysis (Figure 2H). Western blotting to detect phosphorylated S6 and eEF2 confirmed that DGAT1 inhibition led to rapid loss of mTOR and S6K activities in multiple melanoma cell lines with up-regulated expression of endogenous DGAT1, including LOXIMVI and SKMEL5 that harbor *DGAT1* amplification (Figure 3B and Supplementary Figure 3B). This effect was not observed with DGAT2 inhibition (Supplementary Figure 3C). Accordingly, DGAT1 depletion but not DGAT2 depletion using siRNA also resulted in a reduction in phosphorylated S6 and accompanying increase in phosphorylated eEF2 (Figure 3C and Supplementary Figure 3D). Conversely, stable expression of DGAT1 in DGAT1^low^ 888MEL melanoma cells (Figure 1D) as well as transient over-expression of DGAT1 in two other melanoma cell lines resulted in elevated phospho-S6 (Figure 3D and Supplementary Figure 3E).

**Figure 3.**
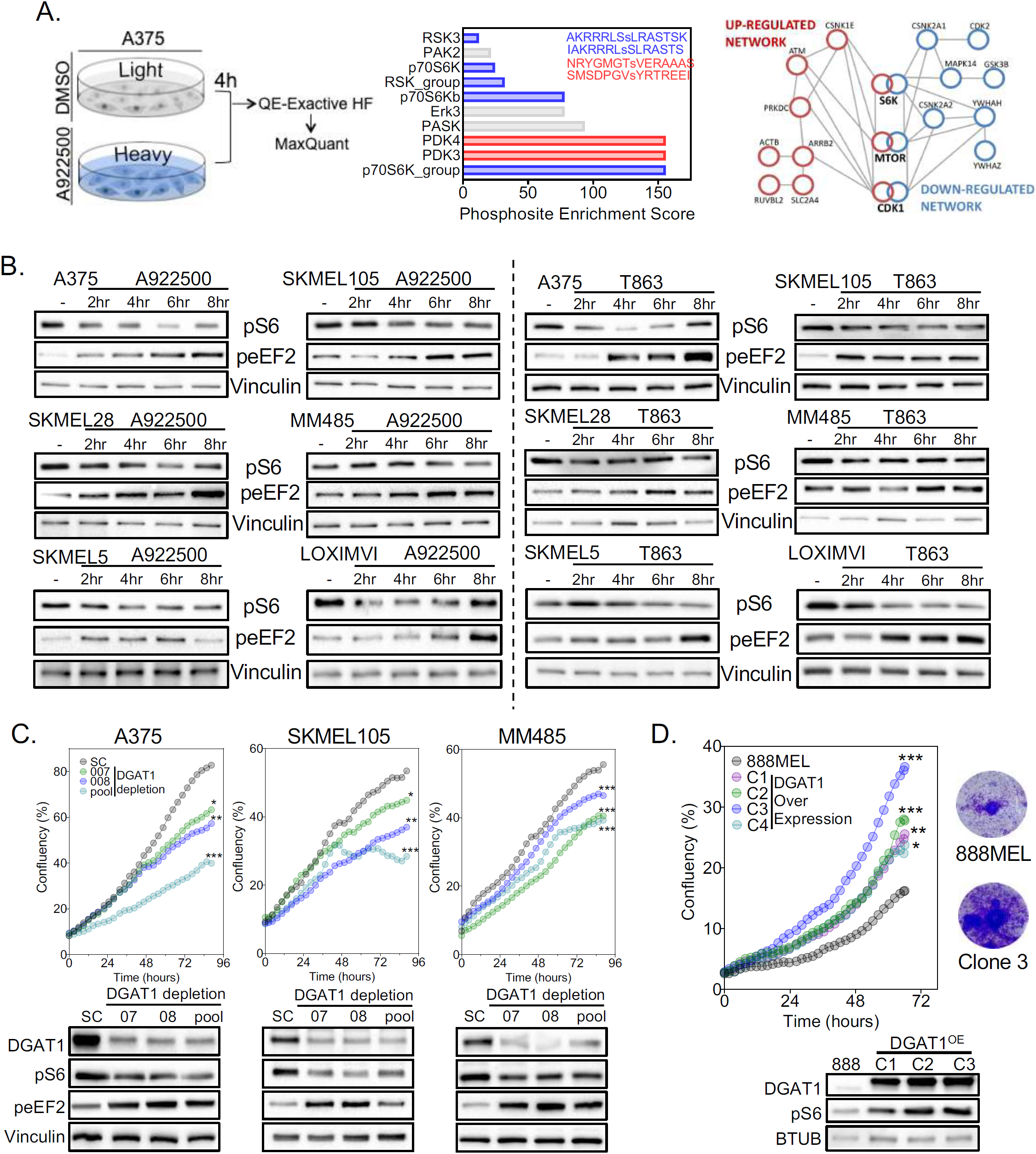
DGAT1 activity is required for maintenance of S6K signalling. (a) Phospho-proteomics work flow (left). Enriched phosphorylated sites (WebGestalt) of up-(59) and down-regulated (96) phosphorylated proteins. Protein hub analysis of both up- and down-regulated phosphorylated proteins (right) (n=3). (b) Protein expression of phospho-S6 and phospho-eEF2 following DGAT1 inhibitor treatment. (c) Confluency curves in cell lines transfected with DGAT1 targeting (007, 008, pool) or scrambled (SC) siRNAs (Mean, n=3) (above). Corresponding protein expression of DGAT1, phospho-S6 and phospho-eEF2. (d) Confluency curves of parental 888MEL cells and 888mel cells following lentiviral transduction with a DGAT1 over-expression vector and clonal selection (Mean, n=3) (upper left). Corresponding crystal violet staining after 72 h growth (upper right). Corresponding protein expression of DGAT1 and phospho-S6 (lower).

Given the effect of DGAT1 antagonism on mTOR signaling, which is heavily implicated in cell growth and division, we next addressed the importance of DGAT1 for melanoma cell proliferation. DGAT1 depletion using siRNA reduced cell proliferation and decreased the fractions of cells in S-phase (Figure 3C and Supplementary Figure 3G), which was in contrast to the increased proliferation and cell cycle progression observed with stable DGAT1 overexpression (Figure 3D and Supplementary Figure 3F). To determine whether these effects were due to the enzymatic function of DGAT1 we utilized four selective DGAT1 inhibitors (AZD3988, AZD7687, A229500 or T863). Pharmacological antagonism of DGAT1 suppressed melanoma cell growth over 96h (Figure 4A), accompanied by decreased cell-cycle progression (Figure 4B). In contrast, no changes in cell growth or cell cycle progression were observed following DGAT2 depletion or inhibition (Supplementary Figure 3B, C). Corroborating the link between DGAT1, S6K activity, and proliferation, over-expression of wild-type p70S6K, or a constitutively active form, partially restored cellular proliferation and phospho-S6 levels in DGAT1-suppressed melanoma cells (Figure 4C and Supplementary Figure 4D). Conversely, stable expression of DGAT1 in 888MEL melanoma enhanced cell proliferation and this was reversed by pharmacological antagonism of S6K using LY2584702 or PF-4708671 (Figure 4E). Taken together, we conclude that DGAT1 promotes melanoma cell proliferation by stimulating S6K activity.

**Figure 4.**
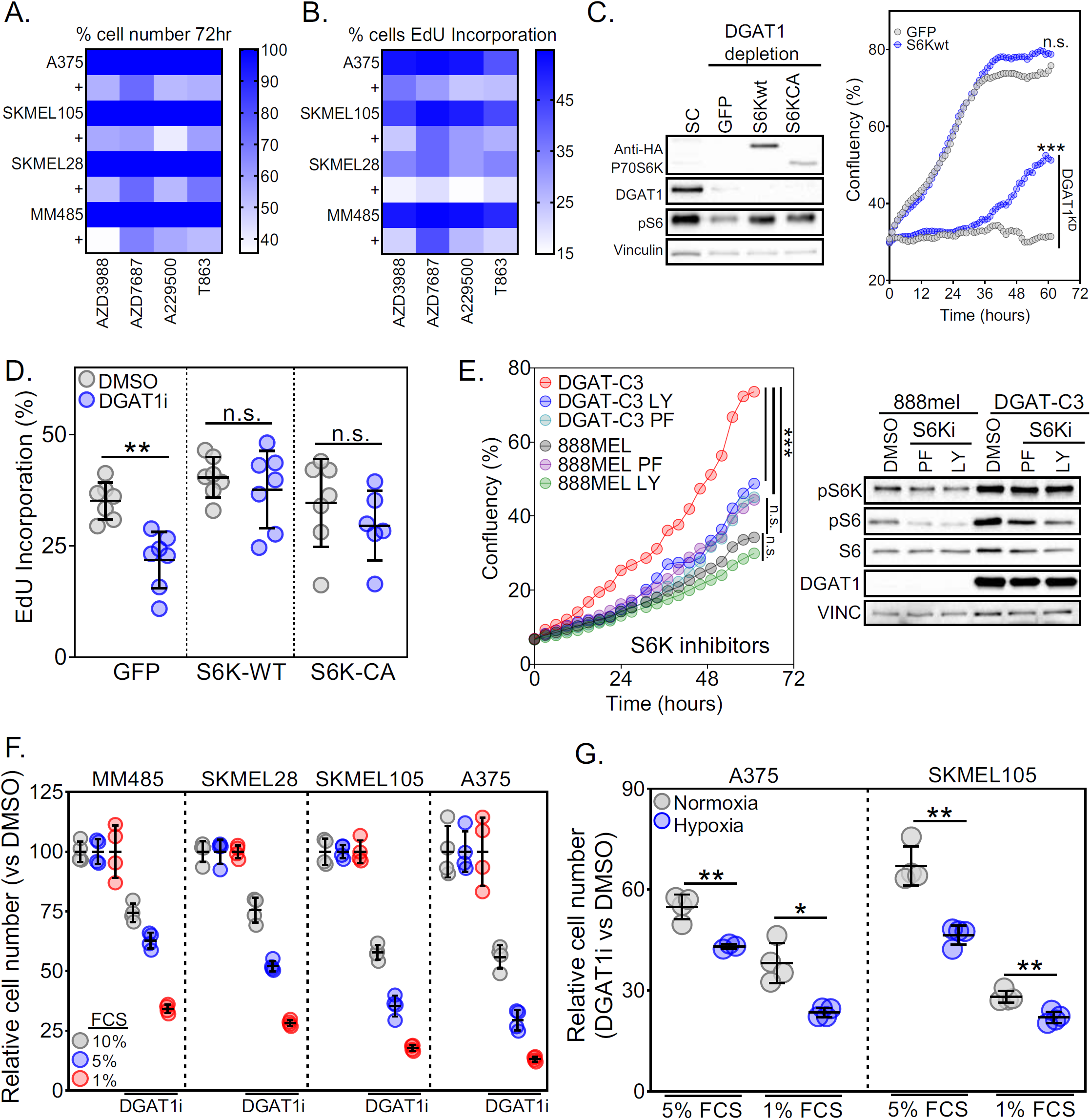
DGAT1 supports S6K dependent melanoma cell growth. (a) Heatmap of relative cell number determined by crystal violet following 72 h DGAT1 inhibitor treatment (Mean, n>3). (b) Percentage of cells in S-phase using EdU incorporation following 24 h DGAT1 inhibitor treatment (Mean, n>3). (c) Protein expression in SKMEL105 cells of HA-tag, DGAT1 and phospho-S6 following both over expression of either GFP wild type S6 kinase or constitutively active S6-kinase and transfection with a DGAT1 targeting or scrambled siRNA (left). Confluency curves of SKMEL105 cells following both over-expression of S6 kinase or GFP and transfection with a DGAT1 targeting or scrambled siRNA (Mean, n=3) (right). (d) Quantification of the percentage of A375 cells in S-phase using EdU incorporation following over-expression of either S6 kinase, constitutively active S6 kinase or a GFP control with/without the presence of A922500 for 24 h (Mean ± SD, n>6). (e) Confluency curves of 888MEL and clone 3 cells following treatment with/without 1 μM LY2584702 or 10 μM PF-4708671 determined by time-lapse microscopy using an Incucyte zoom system (left) (Mean ± SD, n=4). Corresponding protein expression of phospho-P70S6K, phospho-S6, S6 and DGAT1 (right). (f) Graph showing relative cell number determined by crystal violet following 72 h DGAT1 inhibitor treatment with cells grown in varying concentrations of foetal calf serum (FCS) (Mean, n>3). (g) Graph of relative cell number determined by crystal violet following 48 h DGAT1 inhibitor treatment under normoxic or hypoxic conditions (1% O2) with cells grown in varying concentrations of foetal calf serum (FCS) (relative to DMSO control for each condition) (Mean, n>3).

Cancer cells are capable of sustaining cell growth despite transient or limited nutrient availability in the tumor microenvironment arising from poor vascularization (29). Given the connection suggested by gene set enrichment analysis between genes differentially expressed in Dgat1a over-expressing tumors and HIF1 regulation of glycolysis (Figure 2H), we investigated the role of DGAT1 in allowing cancer cells to tolerate nutrient and oxygen deprivation in culture. First, we found that DGAT1 antagonism greatly impaired proliferation when external lipid sources were restricted, with the greatest reduction in cell number observed at the lowest levels of serum (Figure 4F). The suppressive effect of DGAT1 inhibition on S6K activity was also observed in lipid-restricted media (Supplementary Figure 4E). Second, we explored the importance of DGAT1 under hypoxic conditions. Again, we found that DGAT1 inhibition had a profound effect on cell number under low oxygen conditions, which was further exacerbated by limiting serum (Figure 4G). Relative to growth of non-transfected cells, transient DGAT1 overexpression augmented melanoma cell growth to a greater extent under hypoxic conditions (Supplementary Figure 4F). Taken together up-regulation of DGAT1 confers a growth advantage to melanoma cells, which is even more profound under stress conditions likely to be encountered in the tumor microenvironment.

### DGAT1 is essential for lipid droplet formation and acts as a caretaker of mitochondrial health

The capacity of DGAT1 to sequester FA and DAG in LD has the potential to affect both cell growth signaling and ATP production, as FA, DAG and their derivatives are allosteric regulators of various metabolic enzymes, kinases and transcription factors, and FA are also fuel molecules. Therefore, to address directly the effect of DGAT1 on melanoma lipid metabolism using an unbiased approach, we performed Ultra-High-Performance Liquid Chromatography-Mass-Spectrometry (UHPLC-MS) to identify and contrast lipid species extracted from NRAS^mut^ Dgat1a over-expressing or NRAS^mut^ EGFP-expressing zebrafish tumors. This analysis revealed increased concentrations of almost all TAG species detected in tumors with forced Dgat1a expression, with longer poly-unsaturated FA (PUFA) chains showing the greatest increases (Figure 5A, Supplementary Figure 5A and Supplementary Table 3). However, after correction for multiple testing, the changes were no longer significant. This lack of significance does not necessarily disprove the hypothesis that DGAT can impact TAG levels, as several experimental limitations exist that may confound the analysis. First, EGFP-expressing tumors also over-express Dgat1a and have ample LD (19). Second, lipids originating from associated stromal cells dilute the lipids derived from tumor cells. Both issues mean that lipidome differences between Dgat1-over-expressing and EGFP-expressing tumors are likely to be reduced in magnitude.

**Figure 5.**
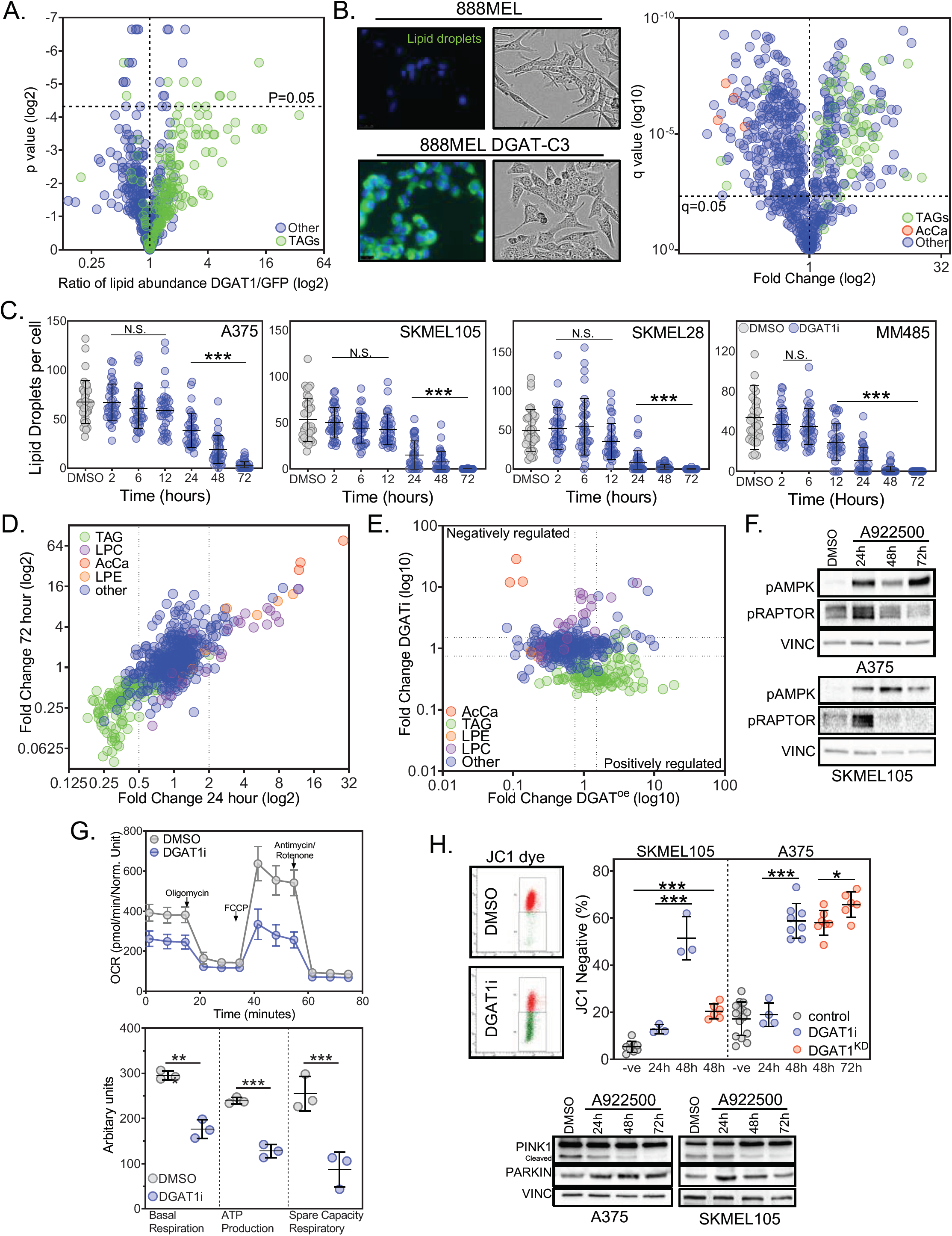
DGAT1-driven lipid droplets act as caretakers of mitochondrial health. (a) Lipidomic profiling using UHPLC-MS of *NRAS*^*G12D*^-positive EGFP-expressing (n=6) and *NRAS*^*G12D*^-positive Dgat1a-over-expressing (n=6) tumors showing the ratio of lipid species (annotated by MS/MS). (b) Representative images of 888MEL and Clone 3 cells stained with BODIPY (left). Brightfield images of 888MEL parental cells and Clone 3 DGAT1 over-expressing cells (middle). UHPLC-lipidomic analysis of 888MEL parental and Clone 3 DGAT1 over-expressing cells. Fold-change relative to 888MEL parental cells (right). TAG (triacyclglycerides), AcCa (acyl carnitine). (c) Quantification of the number of lipid droplets per cell using BODIPY staining following AZD3988 treatment (Mean ± SD, n>30). (d) UHPLC-lipidomic analysis of SKMEL105 cells following A922500 treatment. Fold-change relative to DMSO. TAG (triacyclglycerides), LPC (lysophosphatidycholine), AcCa (acyl carnitine), LPE (lysophosphatidylethanolamine). (e) Lipid species fold changes in SKMEL105 following 72 h A922500 treatment plotted versus lipid species fold changes observed in clone 3 cells. f) Protein expression of phospho-AMPK and phospho-RAPTOR following A922500 treatment. (g) Oxygen consumption rate in A375 cells following 48 h A922500 treatment (upper). Basal respiration, ATP production and spare respiratory capacity were calculated (lower) (Mean ± SD, n=3). (h) Staining with JC-1 dye following A922500 treatment or following transfection with DGAT1 targeting siRNA. The percentage of cells that lost red J-aggregates was calculated by using 1 μM CCP as a positive control and comparing this to untreated cells to create two populations of cells for flow cytometry analysis (upper) (Mean ± SD, n>3). Protein expression of PINK1 and PARKIN following A922500 treatment (lower).

In order to overcome the complexity of whole tumor analysis, we turned to melanoma cell lines. Over-expression of DGAT1 in human melanoma cells resulted in a striking increase in LD (Figure 5B) consistent with cells adopting an engorged morphology (Figure 5B). In parallel, UHPLC-MS lipidomic analysis revealed an increase in almost all TAG species (Figure 5B and Supplementary Table 4, 5), corroborating the trend in TAG seen in zebrafish tumors over-expressing Dgat1a (Figure 5A). Conversely, inhibition of DGAT1 in melanoma cell lines over-expressing endogenous DGAT1 led to a reduction in the amount of LD beginning between 12 and 24h and continuing until 72h (Figure 5C and Supplementary Figure 5B, C), as previously reported for other cell types (30). In contrast, LD were not affected by DGAT2 depletion (Supplementary Figure 5D). As expected, UHPLC-MS lipidomic analysis revealed a reduction of multiple TAG species already at 24h that was maintained until 72h, (Figure 5D, Supplementary Figure 5E and Supplementary Table 6, 7), with TAG containing PUFA particularly affected (Supplementary Figure 5A).

Alongside the anticipated changes in TAG levels following manipulation of DGAT1 activity, we also observed a decrease in several acyl carnitine (AcCa) species upon DGAT1 overexpression (Figure 5B and Supplementary Table 4, 5) and a reciprocal increase in AcCa species following DGAT1 inhibition (Figure 5D, E, Supplementary Figure 5E and Supplementary Table 6, 7). AcCa, formed by esterification of FA (typically long-chain) to L-carnitine, are transported into mitochondria to be processed into acetyl-CoA through β-oxidation. The production of AcCa determines the rate of FAO (31). PDK3 and PDK4 are mitochondrial kinases that suppress conversion of pyruvate to acetyl-CoA when alternative fuel sources to glucose, such as FA, are available (32). The increase in PDK3 and PDK4 activity we detected following DGAT1 inhibition through our phospho-proteomic analysis (Figure 3A) was therefore consistent with increased AcCa availability and hence increased FAO. Moreover, excessive FAO can result in mitochondrial overloading and dysfunction (33). Indeed, we observed evidence of impaired ATP production following DGAT1 inhibition, implied by the increased phosphorylation of both AMPK and RAPTOR (Figure 5F). Further, we observed reduced oxygen consumption in DGAT1 inhibited cells after 48h (Figure 5G), implying decreased mitochondrial respiratory function despite levels of AcCa remaining elevated (Figure 5D and Supplementary Figure 5E). Additionally, post 24h DGAT1 inhibition, we observed loss of mitochondrial membrane potential, decreased PINK1 cleavage, and increased levels of mitophagy factor PARKIN (Figure 5H). Moreover, by blocking AcCa synthesis using Etomoxir (a carnitine palmitoyltransferase inhibitor), we could partially restore mitochondrial membrane potential and cell proliferation (Supplementary Figure 5F, G). Thus, we conclude that DGAT1 maintains mitochondrial function in melanoma cells by regulating availability of FA for FAO.

### DGAT1 promotes survival of melanoma cells in the presence of ROS

To understand the effects of prolonged DGAT1 inhibition (post 72hr) on cellular signaling and metabolism we performed unbiased MS-based whole proteome analysis in SILAC-labelled A375 cells. Analysis of differentially expressed proteins highlighted three key areas: i) FA oxidation (consistent with increased AcCa availability and PDK activity), ii) PPAR signaling presumably indicating the increased availability of lipid regulators of PPAR activity, and iii) NRF2 signaling, which is a well-established response to ROS production that dampens ROS-mediated cellular damage (34) (Figure 6A, and Supplementary Figure 6A-B). Quantitative gene expression analysis further corroborated the effects of DGAT1 inhibition on these three key biological processes (Figure 6B and Supplementary Figure 6C).

**Figure 6.**
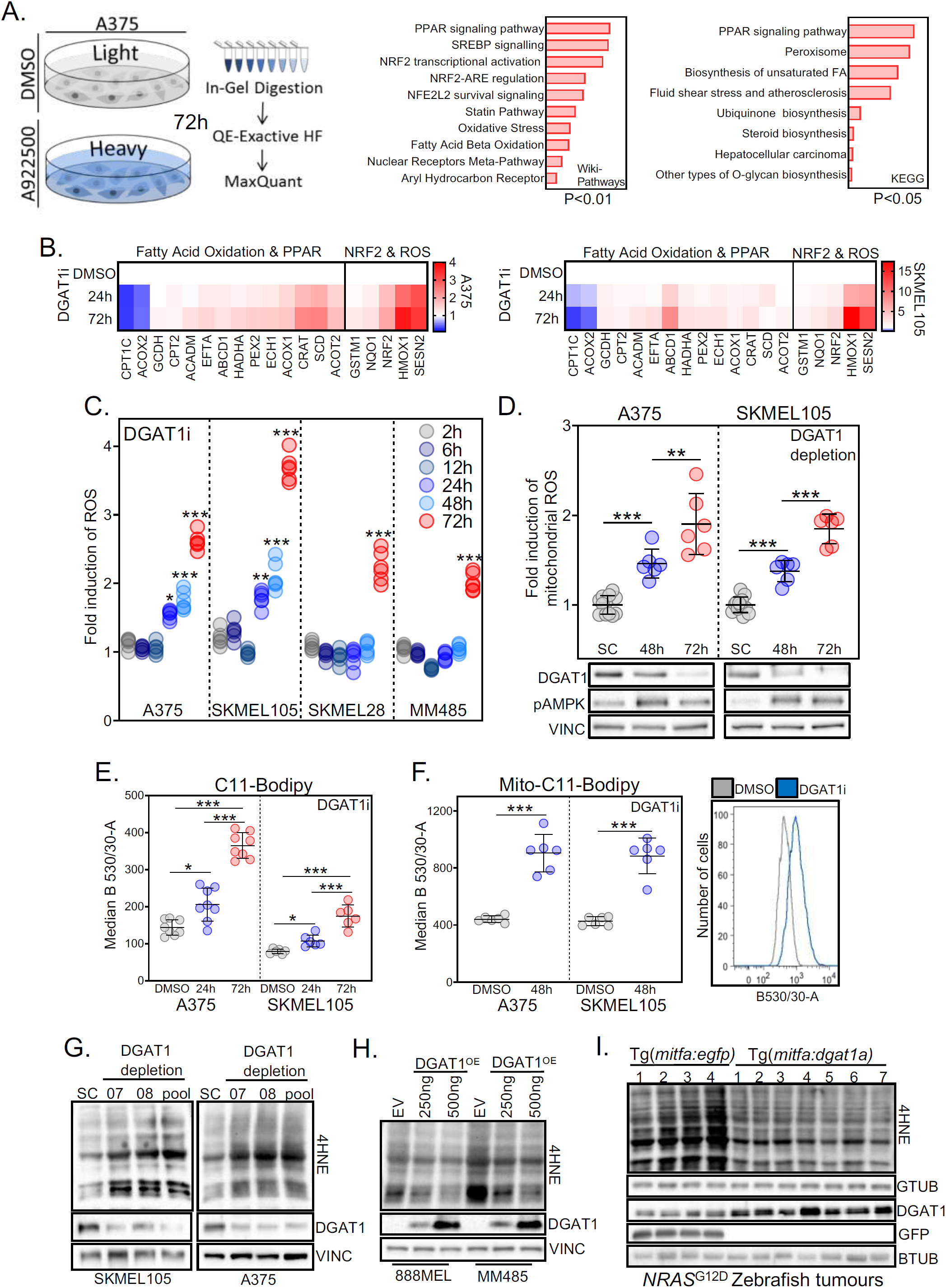
DGAT1 suppression generates ROS leading to mitochondrial lipid peroxidation. (a) Total proteomics work flow (left). GEO of up-regulated proteins (114) ranked by combined score (wikipathways) or log adjusted P values (metascape) (right) (b). RT-qPCR analysis following A922500 treatment. Fold-change relative to DMSO (Mean, n=3). (c) Quantification of ROS levels using dihydroethidium fluorescence following A922500 treatment. Fold-change relative to DMSO (Mean, n>4). (d) Quantification of ROS levels using dihydroethidium fluorescence following transfection with either DGAT1 targeting siRNA or a scrambled control for 48-72 h. Fold-change calculated relative to scrambled control (Mean ± *S*D, n=3) (upper). Corresponding protein expression of DGAT1 and phospho-AMPK (lower). (e) C11-Bodipy staining following A922500 treatment. Median fluorescence determined using FACS (Median ± SD, n>6). (f) Mito-C11-Bodipy staining following A922500 treatment. Median fluorescence determined using FACS (Median ± SD, n>6) (left) Representative histogram of B530/30-A fluorescence (right). (g) Protein expression of DGAT1 and 4-Hydroxynonenal following transfection with either DGAT1 targeting siRNA or a scrambled control for 48 h. (h) Protein expression of DGAT1 and 4-Hydroxynonenal following transfection with either DGAT1 over-expression vector or an empty vector control for 48 h. (i) Protein expression of Dgat1, GFP and 4-Hydroxynonenal in NRAS^G12D^ -positive GFP-expressing (n=4) and NRAS^G12D^ -positive Dgat1a over-expressing (n=7) tumors.

To measure the impact of DGAT1 suppression on ROS more directly, we stained cells with fluorescent probes that are uncaged by ROS. Using this method, we observed increasing levels of ROS over time upon both DGAT1 inhibition and depletion in multiple melanoma cell lines (Figure 6C,D and Supplementary Figure 6D,E), including in the mitochondria specifically (Supplementary Figure 6F), a predicted consequence of excessive FAO (33). Through a chain reaction, oxygen-centered ROS can yield highly reactive cytotoxic lipid species termed lipid peroxides responsible for a form of programmed cell death termed ferroptosis (35). Accordingly, in parallel with increased ROS generation, DGAT1 inhibition led to an increase in the peroxidation of lipids from 24h, with further increases by 48h both in the cytoplasm (Figure 6E) and mitochondria specifically (Figure 6F). Additionally, depletion of DGAT1 resulted in increased protein attachment of 4-Hydroxynonenal (4HNE) (Figure 6G), a by-product of lipid peroxidation. Conversely, we observed decreased protein attachment of 4-Hydroxynonenal (4HNE) in DGAT1 over-expressing melanoma cells (Figure 6H) and also in zebrafish tumors over-expressing Dgat1a (Figure 6I), indicating that lipid peroxidation and its suppression by DGAT1 occurs not only in cell lines but also within tumors.

We next investigated whether the induction of ROS upon DGAT1 suppression affected the survival of melanoma cells. In parallel with ROS generation, we observed induction of ROS response markers, including SOD1 and SOD2 (Figure 7A, Supplementary Figure 7A), and apoptosis induction in some DGAT1^high^ cell lines after prolonged DGAT1 suppression (Figure 7A, B and Supplementary Figure 7B); again, an effect not observed following DGAT2 inhibition (Supplementary Figure 7C, D). Moreover, ROS scavengers Tempol and Ebselen partially suppressed apoptosis induced by DGAT1 depletion (Figure 7B). Conversely, stable over-expression of DGAT1 in DGAT1^low^ 888MEL melanoma cells was protective against ROS-mediated cell death triggered by chemical ROS inducers (Figure 7C). Further underpinning the observation that DGAT1 confers a protective effect on melanoma cells in stressful conditions, DGAT1 over-expression significantly increased cell survival under hypoxic conditions (even with limited serum). Further, SOD2 induction was tempered by elevated DGAT1 activity, indicating suppression of ROS induction (Figure 7D). Thus, DGAT1 promotes the survival of melanoma cells by suppressing ROS production and lipid peroxidation that would otherwise occur as a result of cancer specific altered FA metabolism.

**Figure 7.**
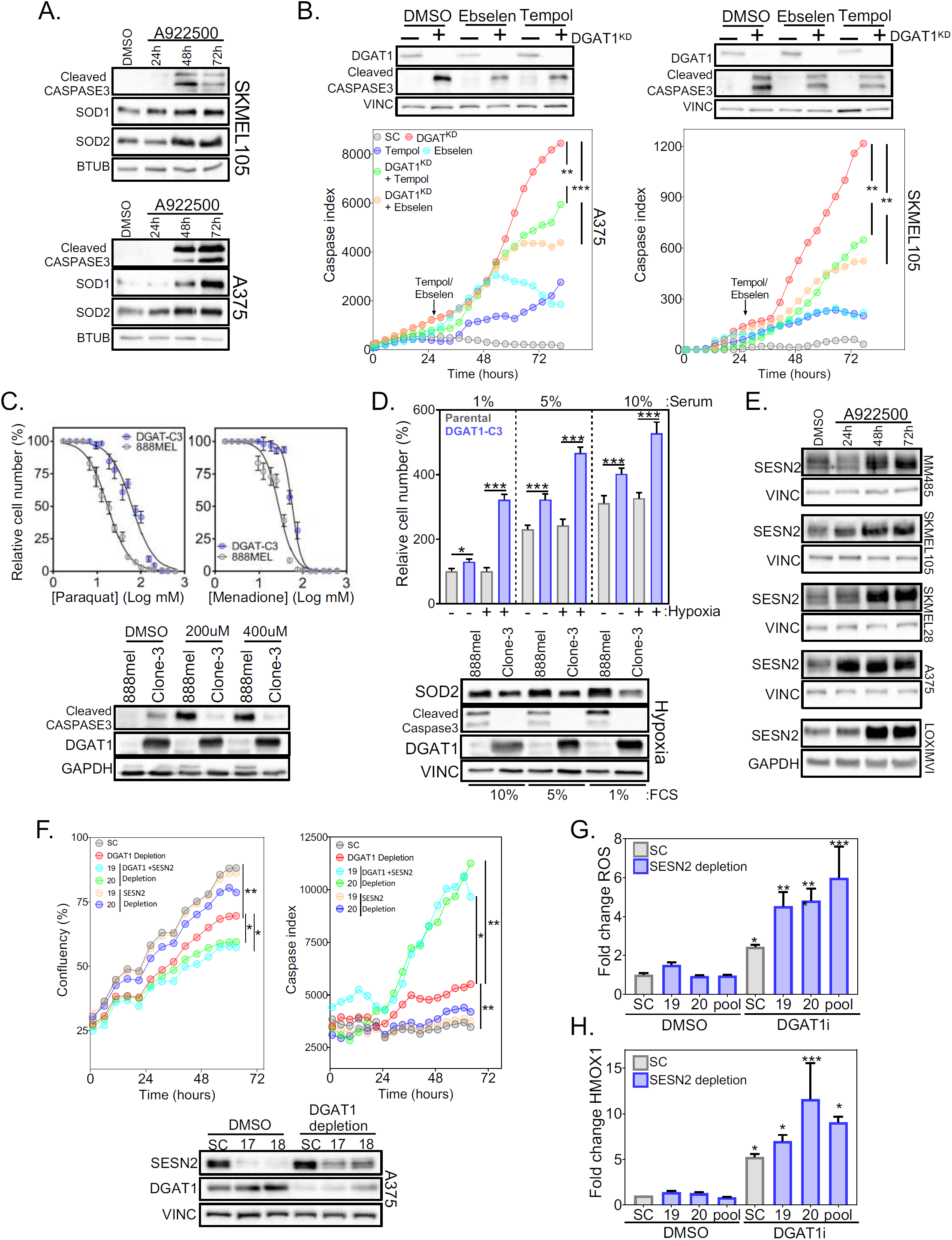
DGAT1 suppression triggers ROS induced apoptosis that is ameliorated by SESN2. (a) Protein expression of SOD1, SOD2 and cleaved caspase3 following A922500 treatment. (b) Cleaved-caspase index following transfection of a DGAT1 targeting or scrambled siRNA. At 24 h cells were treated with/without Tempol or Ebselen (Mean, n=3) (lower). Corresponding protein expression of DGAT1 and cleaved caspase3 (upper). (c) Drug-dose response curve after 72 h treatment with ROS inducers (upper) (Mean ± SD, n*=3*). Protein expression of cleaved-caspase3 and DGAT1 following 72 h paraquat treatment (lower). (d) Relative cell number determined using crystal violet staining following 48 h culture in indicated FCS serum levels under hypoxic (1% O2) or normoxic conditions (Mean ± SEM, n>3) (Upper). Corresponding protein expression of SOD2, DGAT1 and cleaved caspase 3 under hypoxic (1% O2) conditions (lower). (e) Protein expression of SESN2 following A922500 treatment. (f) Confluency curves of A375 cells transfected with either SESN2 or DGAT1 targeting or scrambled siRNA (left). Corresponding cleaved-caspase index (right). Corresponding protein expression of DGAT1 and Sestrin 2 (lower). (g) Quantification of ROS levels using dihydroethidium fluorescence in A375 cells transfected with SESN2 targeting or scrambled siRNA (19, 20) followed by A922500 treatment for 24 h. Fold-change relative to scrambled control (Mean ± SD, n>4). (h) Relative HMOX1 expression in A375 cells transfected with SESN2 targeting or scrambled siRNA (19, 20) followed by A922500 treatment for 24 h. Fold-change relative to DMSO (Mean ± SD, n=3).

DGAT1 inhibition resulted in apoptosis only in a proportion of cells and in a variable fraction dependent on cell line suggesting that not all affected cells produce toxic ROS levels. We therefore hypothesized that anti-ROS mechanisms were activated in some cells, counteracting the effects of DGAT1 suppression. We wanted to investigate how the anti-ROS response was being coordinated downstream of DGAT1 inhibition in order to identify targets that might synergize with DGAT1 inhibition. mRNA for the NRF2 target *SESTRIN2 (SESN2)* was found to be highly up-regulated post DGAT1 inhibition (Figure 6B). Accordingly, SESN2 protein was consistently and significantly up-regulated in a panel of melanoma cell lines following DGAT1 inhibition (Figure 7E). Intriguingly, SESN2 is a major mediator of NRF2 signaling responsible for neutralizing ROS while simultaneously inhibiting mTOR signaling (36). This suggests that SESN2 induction may mediate, but also ameliorate the impact of DGAT1 inhibition on cell growth and survival. Supporting this hypothesis, simultaneous depletion of SESN2 and DGAT1 resulted in a greater reduction in surviving cell numbers (Figure 7F), coinciding with increased apoptosis, compared to DGAT1 depletion or inhibition alone (Figure 7F, Supplementary Figure 7E). Simultaneous knockdown of SESN2 and inhibition of DGAT1 (Supplementary Figure 7F) also led to both elevated levels of ROS (Figure 7G) and increased expression of the NRF2 target gene HMOX1 (Figure 7H) compared to DGAT1 inhibition alone, indicating that without SESN2 induction the neutralization of ROS is impaired. Moreover, we found that the reduction of mTOR signaling induced by DGAT1 inhibition was alleviated by knockdown of SESN2 (Supplementary Figure 7G). Thus, SESN2 acts as a counterfoil to DGAT1, suppressing mTOR signaling while ameliorating ROS inflicted damage.

## DISCUSSION

The role of dysregulated lipid metabolism, encompassing more than just *de novo* FA synthesis, in promoting neoplasia is increasingly apparent (8,9,12,37). However, the adaptations allowing cancer cells to tolerate exposure to excess FA have been overlooked. Using multiple zebrafish models, integrating outputs across multiple omics platforms, including oncogenomic, transcriptomic, proteomic, phospho-proteomic, and lipidomic, combined with assays of cell behavior, we now demonstrate that DGAT1 is a novel *bona fide* metabolism oncoprotein that stimulates melanoma tumorigenesis. We conclude that DGAT1 promotes proliferation through mTOR–S6K signaling. We also show that DGAT1 promotes survival principally by conferring protection against ROS and lipid peroxidation through inducing LD formation (see model; Supplementary Figure 7H).

We uncovered frequent amplification and up-regulation of *DGAT1* in melanoma but also in many other cancers, notably ovarian, breast, uterine, esophageal, liver, pancreatic, head and neck, prostate, stomach and lung cancers. Abundant LD have been observed in a range of cancers, consistent with widespread up-regulation of DGAT1 (38,39), indicating that DGAT1 is likely to perform an oncogenic role in these other cancers too. The co-occurrence of *DGAT1* amplification with *FASN, CD36*, or *MAGL* amplification in human cancers was significant (data not shown), arguing for these co-aberrations being synergistic, in keeping with DGAT1 facilitating safe accumulation of FA in cancer cells.

Multiple indirect mechanisms connect DGAT1 enzymatic activity with S6K activity. On the one hand, by promoting mitochondrial respiratory function and ATP generation, DGAT1 suppresses AMPK thereby relieving inhibition of mTOR (40). Further, through curbing ROS generation by malfunctioning mitochondria, DGAT1 activity represses NRF2 activation and induction of SESN2, a known inhibitor of mTOR (36). Accordingly, knockdown of SESN2 rescued the S6K suppression ensuing DGAT1 inhibition. Other potential DGAT1-dependent factors that could impact mTOR signaling include phosphatidic acid, essential for mTOR activity (41), and DAG, which co-activate PKC that in turn inhibit growth factor signaling to mTOR (42). Levels of these lipids could be rapidly affected by DGAT1-driven sequestration of DAG and FA as TAG in LD. However, changes in concentration of these lipid species following manipulation of DGAT1 activity or expression were not readily apparent from our lipidomics analyses, perhaps limited by their low abundance, instability, rapid association with proteins in the cell, or a combination of all the above. Intriguingly, mTOR–S6K signaling has been shown to induce lipogenesis and LD formation (37,43,44), suggesting that mTOR signaling and lipogenesis are mutually re-enforcing events.

We propose that sequestration of FA as TAG in LD allows cancer cells to accumulate and utilize FA safely, thereby avoiding cell death. As previously described in non-transformed embryonic fibroblasts (30), we observed that DGAT1 suppression resulted in overloading of mitochondria with AcCa, driving excessive FAO and ROS production. Thus, the pathophysiological role of DGAT1 is to regulate the supply of FA to mitochondria within safe limits. Evidently, cancer cells, having elevated FA, require more DGAT1 to achieve this. Lipid peroxides are especially cytotoxic and are generated by the action of oxygen-centered radicals on polyunsaturated fatty acids (PUFA) preferentially (45). As PUFA are concentrated in LD where they are shielded from peroxidation (46), enhanced LD levels mediated by DGAT1 up-regulation confer double protection from cell death by both moderating FAO-triggered ROS production and simultaneously shielding them from PUFA peroxidation.

LD are known to protect non-transformed and transformed cells against a range of cellular stresses typically encountered in the tumor microenvironment, including nutrient deprivation and hypoxia (38,39). Indeed, these conditions induce LD formation downstream of autophagy (30). Additionally, low pH up-regulates DGAT1 and consequently increases LD formation. In turn, LD were required for acidosis-induced metastasis (47). Nutrient deprivation and hypoxia compromise *de novo* FA synthesis and desaturation. Unsaturated FA are essential for membrane fluidity and cell viability, and in the absence of desaturation reactions, or the ability to scavenge unsaturated FA from the circulation, cancer cells must draw on LD for unsaturated FA in order to survive (48). The above role for LD is consistent with our observation that the effect of DGAT1 suppression on cell number was exacerbated by serum withdrawal and hypoxia and explains why conversely DGAT1 overexpression conferred its greatest growth advantage under these conditions.

Our data indicate that DGAT1 inhibition alone could be therapeutic in tumors addicted to DGAT1, although diarrhea attendant on blocking fat absorption through the gut (49) would have to be managed in patients. The identification of signaling molecules such as SESN2, whose suppression synergized with DGAT1 antagonism to produce high levels of ROS and significantly elevated cell death, might one-day feed into a strategy that allows DGAT1 inhibitor concentration to be reduced to a well-tolerated dose. Moreover, such pro-oxidant therapies could inhibit the progression of melanoma, as oxidative stress limits melanoma metastasis (50).

Up-regulation of DGAT1 likely has additional profound consequences for cancer progression, perhaps explaining the association between *DGAT1* amplification and progression-free survival that we uncovered. Enhanced FA uptake and FAO are particularly prominent in tumor initiating cells (9) and DGAT1 could play a critically important role in this crucial subset of tumor cells. It was recently shown that FAO activity correlates with antigen presentation in melanoma and responsiveness to immune checkpoint immunotherapy (51). In fatty liver disease, lipid peroxide adducts were shown to be highly immunogenic and drive adaptive immune responses (52). We found that both substrate production for FAO and lipid peroxidation were suppressed by DGAT1 up-regulation. Moreover, it has been found that LD are enriched with COX2 and arachidonic acid and are major sites of PGE_2_ synthesis (53). PGE_2_ has wide ranging suppressive effects on cellular immunity (54). Thus, DGAT1 up-regulation could promote immune evasion by tumor cells and potentially diminish immunotherapy responses; DGAT1 inhibitors might therefore synergize with immunotherapy.

Overall, our findings demonstrate that overexpression of the oncoprotein DGAT1, with or without gene amplification, is a beneficial adaptation of cancer cells. DGAT1 overexpression permits accumulation of fatty acids, useful as an energy source, for lipogenesis and for signal transduction, while avoiding suppression of growth factor signaling and induction of oxidative stress.

## ACKNOWLEDGEMENTS

We thank David Knight and the staff of the Biological Mass Spectrometry (BioMS) facility, Michael Jackson in the FACs facility, staff of the Bio-imaging facility, The Genomic Technologies Core Facility (GTCF) (all FBMH, UoM) for experimental support. We thank Olivia Sloss and Angeliki Malliri for critically reading the manuscript. AH was funded by grant ERC-2011-StG-282059 from the ERC, grant 610262 from the Melanoma Research Alliance, and a PhD studentship A23251 from Cancer Research UK to support DJW. CF was supported by the Wellcome Trust (Sir Henry Dale fellowship grant number 107636/Z/15/Z) and Biotechnology and Biological Sciences Research Council (grant number BB/R015864/1). The Medical Research Council (MRC) funded the Phenome Centre Birmingham (MR/M009157/1).

## Author Contributions

Cell line-based experimentation: DJW, MPS, RO, JH. Zebrafish models: APB, DJW, HJ, MG, CC. Proteomics: MPS, DJW, PF, CF. Lipidomics: DJW, MPS, ADS, DR, AJ, GRL, WBD. Bio-informatics analysis: DJW, SO, JW, HF, CF, ADS, AJ, GRL, DR, WBD, MPS. Experimental design: DJW, APB, CF, MPS, AH. Sample acquisition and cell lines: PL, CW, MPS, AH. Conceptualization: APB, MPS, AH. Drafting manuscript: DJW, APB, MPS, AH. Revisions by all authors. Acquisition of funds: CW, CF, AH. Study supervision MPS, AH.

## Competing interest

No competing interests to declare.

**Supplementary Materials** contains supplemental methods, figures and tables.

## Methods

### Zebrafish models

Regulated procedures involving zebrafish were ethically approved by The University of Manchester Animal Welfare and Ethical Review Body (AWERB), or by the UMMS Institution Animal Care and Use Committee (A-2016, A-2171), and carried out under a licence issued by the appropriate national regulatory authority. Zebrafish were housed at ∼28 °C under a 14 h light/10 h dark cycle. Transgenic zebrafish expressing *BRAF*^*V600E*^ or *NRAS*^*G12D*^ have been previously described (28,55) and were crossed onto a *mitfa*^w2/w2^ (*mitfa*^-/-^) background to suppress melanocyte development and further onto a *tp53*^M214K/M214K^ background to promote tumorigenesis. Melanocyte restoration and simultaneous over-expression of Dgat1a, Dgat2 or EGFP was then achieved by injection of embryos with a mitfa-minigene containing plasmid as previously described (28). Briefly, zebrafish *dgat1a* and *dgat2* were amplified from cDNA of wild-type 48 h post-fertilisation zebrafish embryos, and subcloned into the pDONR221 vector (see Supplementary Table 8 for oligonucleotide sequences). The pDest-mitfa:dgat1a-pA and pDest-mitfa:dgat2-pA destination vectors were created using an LR clonase reaction consisting of p5E-mitfap, pME-dgat1a or pME-dgat2, p3E-pA and an empty destination vector. Expression plasmid was injected into zebrafish zygotes along with Tol2 mRNA. pCS2-TP plasmid for Tol2 mRNA generation was a kind gift from Dr Koichi Kawakami (National Institute of Genetics). Sufficient embryos for all experimental arms were generated simultaneously, pooled and then randomly assigned to a construct, although formal randomization techniques were not used. Zebrafish were group-housed according to the construct. Only zebrafish embryos with near complete melanocyte rescue at 5 days were retained for further analysis. Analysis of tumor formation was not performed blinded to the construct identity. Sample sizes were not predetermined based on statistical power calculations but were based on our experience with these assays. To assess the statistical significance of differences in overall survival, we used Mantel– Cox’s log-rank tests.

### Cell Lines

Human melanoma cell lines were cultured in High Glucose DMEM with 10 % FBS, and penicillin–streptomycin (Sigma) at 37 °C and 5 % CO_2_. Normal human melanocytes were purchased from Cascade Biologics and cultured according to manufacturer’s guidelines. Lenti-X cells were cultured in High Glucose DMEM with 10 %v/v FBS, and penicillin–streptomycin (Sigma) at 37 °C and 5 % CO_2_. All cells tested negative for mycoplasma and cell lines were authenticated using STR profiling.

| Cell line | Genetic drivers | DGAT1 status |
| --- | --- | --- |
| 888MEL | BRAF<sup>V600E</sup> | DGAT1 low |
| SKMEL28 | BRAF<sup>V600E</sup> | DGAT1 medium |
| SKMEL2 | NRAS<sup>Q61R</sup> | DGAT1 over expressed |
| MM485 | NRAS<sup>Q61R</sup> | DGAT1 over expressed |
| A375 | BRAF<sup>V600E</sup> | DGAT1 over expressed |
| SKMEL105 | BRAF<sup>V600E</sup> | DGAT1 over expressed |
| LOXIMVI | BRAF<sup>V600E</sup> | DGAT1 amplified |
| SKMEL5 | BRAF<sup>V600E</sup> | DGAT1 amplified |

### Compounds and Antibodies

Compounds were used at the following concentrations unless otherwise noted 50 μM AZD3988 (Tocris), 30 μM A922500 (Stratech) 50 μM AZD7687, 70 μM T863 (Sigma), 1 μM Oligomycin (Sigma), 0.5 μM FCCP (Sigma), 1 μM Antimycin-A (Sigma), 1 μM Rotenone (Sigma), 50 μM PF-06424439 (Sigma), 5 μM Ebselen (Tocris), 1 mM Tempol, 200 μM Paraquat (Sigma), Menadione (Sigma), 100 μM Etomoxir (Sigma), 1 μM LY2584702 (Stratech), 10 μM PF-4708671 (Generon).

Antibodies against DGAT1 (ab54037), phospho-PDE1a (ab92696) and 4-Hydroxynonenal (ab46545) were purchased from abcam. Antibodies against Vinculin (66305-1-Ig), Beta-Tubulin (10094-1-AP), PINK1 (23274-1-AP), Parkin (14060-1-AP), SOD1 (10269-1-AP), SOD2 (24127-1-AP), Sestrin 2 (10795-1-AP), GAPDH (60004-1-Ig), and PDK4 (12949-1-AP) were purchased from Proteintech. Antibodies against phospho-S6 (2215), phospo-eEF2 (2331), phospho-AMPK (50081), phospho-RAPTOR (2083), Caspase-3 (9662), phospho-P70 S6 kinase (9206), P70 S6 Kinase (2708), S6 (2317) and GFP (2956) were purchased from Cell Signalling. The Antibody against HA (901533) was purchased from Biolegend. The Antibody against gamma-tubulin (T5326) was purchased from Sigma.

### Plasmids & siRNA

All plasmids were transfected using Lipofectamine (Invitrogen) following standard protocols. The plasmids used were purchased from Addgene: pRK7-HA-S6K1-WT (8984); pRK7-HA-S6K1-F5A-E389-deltaCT (8990); pcDNA3.1-mMaroon1 (83840). The GFP and WPRE elements were excised from pCDH-MCS-T2A-copGFP (a kind gift from Andrew Gilmore, The University of Manchester) using BspEI and KpnI. mApple (BspEI and XhoI adapters) and WPRE (XhoI-KpnI adapters) were PCR amplified, digested and subcloned to create the pCDH-MCS-T2A-mApple vector (see Supplementary Table 8 for oligonucleotide sequences). DGAT1 was further subcloned into the both the pCDNA3.1 vector and pCDH-MCS-T2A-mApple using the MCS.

All siRNA was transfected using Lipofectamine RNAi Max (Invitrogen) following standard protocols. The following siRNA were ordered from Dharmacon: DGAT1 007 5’ UCAAGGACAUGGACUACUC 3’; DGAT1 008 5’ GCUGUGGUCUUACUGGUUG 3’; DGAT1 smart pool #J-009333-00-0005; DGAT2 01 5’ GAACACACCCAAGAAAGGU 3’; DGAT2 02 5’GGAGGUAUCUGCCCUGUCA3’; DGAT2 03 5’ UCAUGGAGCUGACCUGGUU 3’; DGAT2 04 5’GAAUGCCUGUGUUGAGGGA 3’; DGAT2 smart pool #J-009333-08; SESN2 19 5’GGAGGGAGUAUUAGAUUAU3’; SESN2 20 5’GCAGGGACCCGUUGAACAA3’. The scrambled control siRNA (SIC002) was ordered from Sigma.

### Viral transduction

Briefly, Lenti-X cells were transfected with pMDLg/pRRE, pMD2.G, pRSV-Rev plasmids (all kind gifts from Angeliki Malliri, Cancer Research UK Manchester Institute) and pCDH-EF1α-DGAT1-T2A-mApple viral vectors using Fugene (Promega) following standard protocols. The viral containing supernatant was filtered using a 0.45 μm filter and frozen at –80 °C prior to transduction of target cells. The supernatant containing the viral particles was added to target cells along with 10 ng/ml Polybrene (Millipore) for 24 h. Target cells were then grown and selected from single cell colonies.

### Protein lysate preparation and Western Blotting

Cells were washed with PBS and lysed with sample buffer (62.5 mM TRIS pH 6.8, 2 %w/v Sodium dodecyl sulfate (SDS), 10 %v/v glycerol, 0.01%w/v bromophenol blue, 3 %v/v 2-mercaptoethanol). Lysates were then sonicated and heated to 95 °C for 10 minutes prior to being evenly loaded onto SDS-polyacrylamide gels using the Mini Trans-Bot electrophoresis system (Biorad), followed by transfer to PVDF using standard western blotting procedures.

### Lipid droplet staining and image analysis

#### Bodipy 493/503

Indicated cells were stained with 2 μM Bodipy 493/503 (ThermoFisher Scientific) and 5 ng/ml Hoecsht 3342 (Cell Signalling) for 30 minutes prior to fixing in 4 %w/v paraformaldehyde and imaging using a Leica microscope system. Images were processing using Fiji.

#### Lipidtox

Indicated cells were fixed in 4 %w/v paraformaldehyde and stained with LipidTox Green (ThermoFisher Scientific) according to manufacturer’s instructions, and 5ng/ml Hoecsht 3342 (Cell Signalling) for 15 minutes prior to imaging using a Leica microscope system. Images were processing using Fiji.

### RNA Isolation and real-time PCR analysis

RNA from cell lines was isolated with TRIZOL® (Invitrogen). After chloroform extraction and centrifugation, 5 µg RNA was DNase treated using RNase-Free DNase Set (Qiagen). 1 µg of DNase treated RNA was then taken for cDNA synthesis using the Protoscript I first strand cDNA synthesis kit (New England Biolabs). Selected genes were amplified by quantitative real time PCR (RT-qPCR) using Sygreen (PCR Biosystems). Relative expression was calculated using the delta-delta CT methodology and beta-actin was used as reference housekeeping gene. Sequences for primers used can be found in the Supplementary Table 8.

### Incucyte cell-proliferation assay and apoptosis assay

Indicated cell lines were seeded into 24-well plates at a density of 15,000–20,000 cells per well, depending on growth rate and the design of the experiment. After 24 h drugs or siRNA were added and cells were imaged every hour using the Incucyte ZOOM (Essen Bioscience) Phase-contrast images were analysed to detect cell proliferation based on cell confluence. For cell apoptosis, caspase-3 and caspase-7 green apoptosis-assay reagent (Life Technologies) was added to the culture medium following manufacturer’s instructions. Cell apoptosis was analysed based on green fluorescent staining of apoptotic cells.

### Flow cytometry

#### Mitochondrial Membrane potential

Indicated cell lines were trypzinized and pelleted by centrifugation at 500 g for 5 min, washed with PBS. For mitochondrial membrane potential cells were stained with 2 μM JC-1 (Life Technologies) for 30 minutes at 37 °C. For positive control samples 0.5µM FCCP was added simultaneously with JC-1. Data was acquired by the BD BIOsciences Foretessa and quantified using the Flowjo software. A minimum of 10,000 cells were analysed per condition.

#### Lipid peroxidation

Indicated cell lines were trypzinized and pelleted by centrifugation at 500 g for 5 min, followed by a PBS wash. For lipid peroxidation cells were stained with either 5 μM BODIPY™ 581/591 C11 (ThermoFisher Scientific) or MitoPerOx (Abcam) for 30 minutes at 37 °C. Data was acquired by the BD BIOsciences Foretessa and quantified using the Flowjo software. A minimum of 10,000 cells were analysed per condition.

#### Mitochondrial ROS

Indicated cell lines were trypzinized and pelleted by centrifugation at 500 g for 5 min, followed by a PBS wash. For mitochondrial specific ROS detection, cells were stained with 2.5 μM Mitosox (ThermoFisher Scientific) for 30 minutes at 37 °C. Data was acquired by the BD BIOsciences Foretessa and quantified using the Flowjo software. A minimum of 10,000 cells were analysed per condition.

### Proliferation Assays

#### Crystal Violet

Indicated cells were stained and fixed with 0.5 %w/v crystal violet (Sigma) in 4 %w/v paraformaldehyde/PBS for 30 minutes. Fixed cells were then solubilised in 2 %w/v SDS/PBS and absorbance measured at 595 nm using Synergy H1 microplate reader (BioTek).

#### EdU Incorporation

Indicated cells were labelled with 20 µM 5-ethynyl-2’-deoxyuridine (EdU) for 4 h and processed following the manufacturer’s protocol (Click-iT® EdU Alexa Fluor® 488 Imaging Kit, Thermo Fisher). Prior to imaging cells were then stained with 5ng/ml Hoecsht 3342 for 15 minutes. Stained cells were analysed using a using a Leica microscope system. Images were processing using Fiji.

### Dihydroethidium Assay

Cells were stained with 5 μM Dihydroethidium for 20 minutes in the dark at 37 °C. Fluorescence was measured at excitation 480nm emission 570 nm using Synergy H1 micro plate reader (BioTek). Fluorescence values were normalised to cell number by staining the cells with crystal violet after fluorescence read.

### Cancer bioinformatics

We evaluated both point mutations and CNV in the TCGA SKCM firehose legacy, TCGA pan-cancer and Cancer Cell Line Encyclopedia datasets using the cBioPortal platform (56). The GISTIC2.0 algorithm was used to identify focal amplifications (57). Gene Ontology analysis was carried out using both enrichR (58) and metascape software (59). Association between mRNA expression in TCGA datasets and survival was evaluated using OncoLnc (60). mRNA levels determined by microarray were accessed through the Oncomine platform (61).

## Quantification and Statistical Analysis

Data was tested for normality using the Shapiro-Wilk test. Data was considered to be normally distributed if p>0.05. Differences in the number of lipid droplets per cell, relative cell number and percentage EdU incorporation between DMSO and drug treated cells were assessed using an unpaired two-sided *t-*test, or Mann-Whitney test if data were not normally distributed. In comparing the differences in these same characteristics between cells transfected with either non-target or one of several siRNA oligonucleotides, a one-way ANOVA with Tukey’s multiple comparisons test (or Friedman with Dunn’s multiple comparisons test if data were not normally distributed) was used to measure significance. Differences were considered significant if p<0.05. All data obtained was analysed using Graphpad Prism 8.1.

## Data and Code Availability

The zebrafish tumor RNA-seq data has been deposited with the Gene Expression Omnibus (GEO) with the accession code GSE144555. The mass spectrometry proteomics data have been deposited with the ProteomeXchange Consortium via the PRIDE partner repository with the dataset identifier PXD017487. Extended Lipidomics Data for zebrafish tumors and human melanoma cell lines can be found in Supplementary Tables 3-7. Code for proteomic analysis can be found at https://github.com/JoWatson2011/DGAT_2019.

