## Supplemental material for "DGAT1 is a lipid metabolism oncoprotein that enables cancer cells to accumulate fatty acid while avoiding lipotoxicity"

Wilcock et al.

**Supplementary Methods**

**Supplementary Tables 1–8**

**Supplementary Figures 1–7**

### Supplementary Methods

#### RNA-seq and Gene Ontology Analysis

Zebrafish tumors were excised and the RNA isolated using RNeasy RNA extraction kit (Qiagen) after homogenisation. RNA-seq libraries were prepared using a TruSeq stranded mRNA sample prep kit and run on a HiSeq 4000 (Illumina) platform. Adapters were trimmed from raw sequencing reads using Trimmomatics v0.32 (1). Trimmed reads were aligned to the zebrafish genome (Ensembl, GRCz11) using STAR v2.5.3 (2) and the Ensembl GRCz11 annotation file. Reads that mapped to chromosomes 1-25 were retained. Gene counts were determined using featureCounts v1.6.2 (3) and differential expression analysis was performed using DESeq2 v1.14.1 (4), using a adjusted p-value cut off of <0.05. DESeq2 was used to generate log2-normalised variance stabilising transformed (VST) counts. Heatmaps were produced using morpheus (https://software.broadinstitute.org/morpheus). Hierarchical clustering was performed using the one minus Pearson correlation method. Prior to Gene Ontology analysis, zebrafish genes were converted to their human orthologue (bioDBnet), the analysis was carried out using enrichR (5) and metascape (6).

#### Proteomics

***SILAC labelling***

For quantitative mass spectrometry, A375 cells were labelled in SILAC DMEM supplemented with 10 %v/v dialyzed fetal bovine serum (Sigma), 2 mM glutamine, 100 U/ml penicillin and 100 µg/ml streptomycin for 15 days to ensure complete incorporation of amino acids (data not shown). Two cell populations were obtained: one labelled with natural variants of the amino acids (light label; Lys0, Arg0) and a second one with heavy variants of the amino acids (L-[13C6,15N4]Arg (+10) and L-[13C6,15N2]Lys (+8)) (Lys8,Arg10). The light amino acids were from Sigma, while their heavy variants were from Cambridge Isotope Labs.

***Sample Preparation for Mass Spectrometry Analysis***

Cells from the two SILAC conditions treated as indicated were lysed at 4°C in ice cold modified RIPA buffer (50 mM Tris, pH 7.5, 150 mM NaCl, 1 %v/v NP-40, 0.1 %w/v sodium deoxycholate, 1 mM EDTA, 5 mM β-glycerolphosphate, 5 mM sodium fluoride, 1 mM sodium orthovanadate, 1 complete inhibitor cocktail Tablet per 50 ml). Proteins were precipitated for two hours at -20°C in four-fold excess of ice-cold acetone. The acetone-precipitated proteins were solubilized in denaturation buffer (10 mM HEPES, pH 8.0, 6 M urea, 2 M thiourea) and the SILAC-labelled lysates were mixed 1:1 based on protein concentrations. Proteins were reduced with 1 mM dithiothreitol (DTT) for 60 min, alkylated with 5.5 mM chloroacetamide (CAA) for 60 min and digested first with endoproteinase Lys-C (Wako, Osaka, Japan) and then, after a five-fold dilution with 50 mM ammonium bicarbonate (ABC), with trypsin (modified sequencing grade, Sigma). The peptide mixture was desalted and concentrated on a C18-SepPak cartridge (Waters, USA) and eluted with 50 %v/v acetonitrile. Phosphorylated peptides were enriched using TiO2beads (5 μm, GL Sciences Inc., Tokyo, Japan), as previously described (7). Equal amounts of SILAC lysates were then mixed 1:1, reduced with DTT, and alkylated with CAA before being resolved on SDS-PAGE (8-12 %w/v, Invitrogen). Separated proteins were fixed in the gel and visualized with colloidal Coomassie staining (Invitrogen). Each gel lane was excised and separated into eight segments that were sliced, destained with 50 %v/v EtOH in 25 mM ABC, and dehydrated with 100 %v/v EtOH. Proteins were digested with sequence-grade trypsin (Sigma) overnight. Trypsin activity was quenched by acidification with trifluoroacetic acid (TFA) and peptides were extracted from the gel sections with increasing concentrations of acetonitrile. Organic solvent was evaporated in a vacuum centrifuge, as described (8).

***Mass Spectrometry Analysis***

Both enriched phosphorylated peptides and in-gel digested peptides were desalted and concentrated on STAGE-tips with two C18 filters and eluted using 40 %v/v acetonitrile, dried and reconstituted in 5 %v/v acetonitrile in 0.1 %v/v formic acid prior to analysis by LC-MS/MS using an UltiMate 3000 Rapid Separation LC (RSLC,Dionex Corporation, Sunnyvale, CA) coupled to a QE HF (Thermo Fisher Scientific, Waltham, MA) mass spectrometer. Mobile phase A was 0.1 %v/v formic acid in water and mobile phase B was 0.1 %v/v formic acid in acetonitrile and the analytical column utilized was a 75 mm x 250 μm inner diameter 1.7 μm CSH C18 (Waters). Samples were transferred to a 5 μl loop before loading on to the column at a flow of 300 nl/min for 5 minutes at 5 %v/v B. The loop was subsequently taken out of line and the peptides separated using a gradient that went from 5 %v/v to 7 %v/v B and from 300 nl/min to 200 nl/min in 1 min followed by a shallow gradient from 7 %v/v to 18 %v/v B in 64 min, then from 18 %v/v to 27 %v/v B in 8 min and finally from 27 %v/v B to 60 %v/v B in 1 min. The column was washed at 60 %v/v B for 3 min before re-equilibration to 5 %v/v B in 1 min. At 85 min, the flow was increased to 300 nl/min until the end of the run at 90 min. Mass spectrometry data was acquired in a data dependent manner for 90 min in positive mode. Peptides were selected for fragmentation automatically by data dependent analysis on a basis of the top 8 (phospho-proteome) or top 12 (proteome) peptides with m/z between 300 to 1750 Th and a charge state of 2, 3 or 4 with a dynamic exclusion set at 15 sec. The MS Resolution was set at 120,000 with an AGC target of 3e6 and a maximum fill time set at 20 ms. The MS2 Resolution was set to 30,000, with an AGC target of 2e5, a maximum fill time of 45 ms, isolation window of 1.3 Th and a collision energy of 28.

***Data Analysis of quantitative MS data***

Raw data were analyzed with the MaxQuant software suite, version 1.5.6.5, with the integrated Andromeda search engine (9). Proteins were identified by searching the HCD-MS/MS peak lists against a target/decoy version of the human Uniprot database, which consisted of the complete proteome sets and isoforms (2016 release) supplemented with commonly observed contaminants such as porcine trypsin and bovine serum proteins. Tandem mass spectra were initially matched with a mass tolerance of 7 ppm on precursor masses and 0.02 Da or 20 ppm for fragment ions. Cysteine carbamidomethylation was searched as a fixed modification. Protein N-acetylation, oxidized methionine and either deamidation of asparagine and glutamine (proteome analysis) or phosphorylation of serine, threonine, and tyrosine (phosphoproteome analysis) were searched as variable modifications. Labelled lysine and arginine were specified as fixed or variable modification, depending on prior knowledge about the parent ion (MaxQuant SILAC identification). False discovery rate was set to 0.01 for peptides, proteins and modification sites. Minimal peptide length was six amino acids. Only peptides with Andromeda score >40 were included. To pinpoint the actual phosphorylated amino acid residue(s) within all identified phospho-peptide sequences, MaxQuant calculated the localization probabilities of all putative phosphorylation sites using the PTM score algorithm as described (10). Potential contaminants, reverse sequenced peptides and phosphorylation sites with a localisation probability of less than 0.75 (class I) (10) were filtered from the dataset. The remaining data were filtered to remove sites or peptides without quantification in at least two of the three replicates for each time point. The median of the replicates was taken. Sites or peptides with a SILAC ratio of greater than 1.5 were considered up-regulated whilst those with a ratio less than 0.75 were considered down-regulated. Correlation was based on Pearson coefficient and visualized in R. The Phosphopeptide enrichment score was calculated using Webgestalt (11). Gene network visualization was performed using enrichR (5). For the proteome analysis a minimum of three to seven peptide identifications with at least two being uniquely assigned to the particular protein were required. Sequence coverage of the identified proteins was at least 5%. Gene Ontology analysis was carried out using enrichR (5) and metascape (6).

**Lipidomics**

***Sample preparation***

Cells were seeded in 6 well plates at a density of 150,000 cells per well and treated with/without A922500 for 24-72 h. Cells were then washed in PBS and snap frozen on dry ice. All data was normalised to cell number calculated by crystal violet staining in wells that had undergone the exact same experimental conditions. Ice-cold 3:1 propan-2-ol: water (both LC-MS grade, VWR) was added to each well containing washed, frozen cells. The amount of solvent added was normalised to the cell density (ranges from 0.9-1.8 mL). Working quickly on wet ice, cells were scraped into the solvent solution using cell scrapers (Corning). Cell and solvent suspension mix was removed to a 2 mL microfuge tube and freeze thawed twice and vortexed (30 s) to lyse cells, precipitate proteins and solubilise lipids. Samples were centrifuged (20,000 g, 4 °C, 20 min) and the total supernatant was taken and dried in a SpeedVac concentrator (Thermo Fisher). An extract blank sample was created by carrying out the above procedure in the absence of cells. For zebrafish tumors, tissue was dissected and snap frozen in liquid nitrogen and stored frozen until lipid extraction. Frozen samples were weighed (mass ranges 6-18 mg) and then homogenised in 45.7 µL/mg wet tissue mass 75:25 propan-2-ol/water (both LC-MS grade, VWR) using a Precellys24 homogeniser and CK14 homogenisation tubes (both Stretton Scientific, UK). Samples were centrifuged (20,000 g, 4 °C, 20 min) and the 250 µL of the supernatant was taken (equivalent to extraction from a 5.5 mg piece of tissue) and dried in a SpeedVac concentrator (Thermo Fisher). An extract blank sample was created by carrying out the above procedure in the absence of tissue. Prior to UHPLC-MS analysis, samples were re-suspended in ice-cold 3:1 propan-2-ol: water (i) for cell extracts this was normalised to cell density readings (added solvent ranges between 100-200 µL); (ii) for tissue extracts a fixed volume of 100 µL was added. Samples were vortexed (30 s) and centrifuged (20,000-g, 4 °C, 20 min). A fixed volume of supernatant (25 µL) was taken from each sample and mixed by vortexing (30 s) to create a pooled QC. The remainder of the supernatant from each sample was removed into HPLC vials. The pooled QC was divided into multiple HPLC vials. All samples were set on the autosampler at 4 °C for immediate UHPLC-MS analysis.

***UHPLC-MS lipidomics***

The samples were maintained at 4 °C and analysed applying two Ultra-High-Performance Liquid Chromatography-Mass Spectrometry (UHPLC-MS) methods using a Dionex UltiMate 3000 Rapid Separation LC system (Thermo Fisher

Scientific, MA, USA) coupled with a heated electrospray Q Exactive Focus mass spectrometer (Thermo Fisher Scientific, MA, USA). Lipid extracts were analysed on a Hypersil GOLD column (100 x 2.1mm, 1.9 µm; Thermo Fisher Scientific, MA, USA). Mobile phase A consisted of 10 mM ammonium formate and 0.1 %v/v formic acid in 60 %v/v acetonitrile/water and mobile phase B consisted of 10 mM ammonium formate and 0.1 %v/v formic acid in 90 %v/v propan-2-ol/water. Flow rate was set for 0.40 mL.min-1 with the following gradient: t=0.0, 20 %v/v B; t=0.5, 20 %v/v B, t=8.5, 100 %v/v B; t=9.5, 100 %v/v B; t=11.5, 20 %v/v B; t=14.0, 20 %v/v B, all changes were linear with curve = 5. The column temperature was set to 55 °C and the injection volume was 2 µL. Data were acquired in positive and negative ionisation mode separately within the mass range of 150 – 2000 m/z at resolution 70,000 (FWHM at m/z 200). Ion source parameters were set as follows: Sheath gas = 48 arbitrary units, Aux gas = 15 arbitrary units, Sweep gas = 0 arbitrary units, Spray Voltage = 3.2 kV (positive ion) / 2.7 kV (negative ion), Capillary temp. = 380 °C, Aux gas heater temp. = 450 °C. Data dependent MS2 in 'Discovery mode' was used for the MS/MS spectra acquisition using following settings: resolution = 17,500 (FWHM at m/z 200); Isolation width = 3.0 m/z; stepped collision energies (stepped CE) = 20, 40, 100 [positive ion mode] / 40, 60, 130 [negative ion mode]. Spectra were acquired in five different mass ranges: 150 – 510 m/z; 500 – 710 m/z; 700 – 860 m/z; 850 – 1010 m/z; 1000 – 2000 m/z. A Thermo ExactiveTune 2.8 SP1 build 2806 was used as instrument control software in both cases and data were acquired in profile mode. Quality control (QC) samples were analysed as the first ten injections and then every seventh injection with two QC samples at the end of the analytical batch. Two blank samples were analysed, the first as the sixth injection and then the second at the end of each batch.

#### *Mass spectrometry raw metabolomics data processing*

Raw data acquired in each analytical batch were converted from the instrument-specific format to the mzML file format applying the open access ProteoWizard (version 3.0.11417) msconvert tool (12). During this procedure, peak picking and centroiding, were achieved using vendor algorithms. Isotopologue Parameter Optimization (IPO - version 1.0.0, using XCMS - version 1.46.0) (13) was used to perform automatic optimization of XCMS (14) peak picking parameters. For centWave peak picking algorithm following parameters and ranges were used: min_peakwidth (from 2 to 10); max_peakwidth (from 20 to 60); ppm (from 5 to 15); mzdiff (-0.001 to 0.01); snthresh (10); noise (10000); prefilter (3); value_of_prefilter (100); mzCenterFun (wMean); integrate (1); fitgauss (FALSE); verbose.columns (FALSE). Optimised XCMS parameters for raw data files deconvolution were: min_peakwidth (6); max_peakwidth (30); ppm (14); mzdiff (0.001); snthresh (10); noise (100); prefilter (3); value_of_prefilter (100); mzCenterFun (wMean); integrate (1); fitgauss (FALSE); verbose.columns (FALSE). For feature grouping method *density* was used with following: minfrac (0.5); minsamp (1); bw (0.25); mzwid (0.01); max (50); sleep (0). A data matrix of metabolite features (*m/z*-retention time pairs) versus samples was constructed with peak areas provided where the metabolite feature was detected for each sample.

#### *Peak matrix processing*

The data for pooled QC samples were applied to perform QC filtering. The first five QCs for each batch were used to equilibrate the analytical system and therefore subsequently removed from the data before the data was processed and analysed.

The data from the pooled QC samples were used to apply QC filtering. For each metabolite feature detected QC samples 1-8 were removed (i.e. leaving a blank and 2 QCs at the start of each batch) and the relative standard deviation and percentage detection rate were calculated using the remaining QC samples. Blank samples at the start and end of a run were used to remove features from non-biological origins. Any feature with an average QC intensity less than 20 times the average intensity of the blanks was removed. Any samples with >50% missing values were excluded from further analysis. Metabolite features with a RSD > 30 % and present in less than 90 % of the QC samples were deleted from the dataset. Features with a <50% detection rate over all samples were also removed. Prior to statistical analysis, the data was normalised using probabilistic quotient normalisation (PQN) (15). For multivariate analysis missing values were replaced by applying *k* nearest neighbour (kNN) missing value imputation (k = 5) followed by log transformation (16).

***Lipid annotation***

LipidSearch (version 4.2, Thermo Fisher Scientific) was used to annotate peaks based on their MS/MS fragmentation patterns. For lipid annotation, all experimental LC-MS/MS spectra data were searched against a MS/MS lipid library in the LipidSearch software database using the following potential ion forms: positive ion = $[M+H]^+$, $[M+NH_4]^+$, $[M+Na]^+$, $[M+K]^+$, $[M+2H]^{2+}$; negative ion = $[M-H]^-$, $[M+HCOO]^-$, $[M+CH_3COO]^-$, $[M+Cl]^-$, $[M-2H]^{2-}$. The quality of the annotation was graded as A-D. This is defined as: Grade A = all fatty acyl chains and class were completely identified; Grade B = some fatty acyl chains and the class were identified; Grade C = either the lipid class or some fatty acyls were identified; Grade D = identification of less specific fragment ions. Only peaks with an MS/MS identification were discussed in this manuscript. All lipid annotations are reported at a confidence of level 3 according to the Metabolomics Standards Initiative (17).

***Univariate Statistics***

For univariate statistics the normalised data was used to avoid including imputed values in the calculations. Fold changes were computed between all pairs of groups. For 2-group comparisons a t-test was applied to determine features showing a significant different between groups. For comparisons exploring two factors a 2-way ANOVA with an interaction term was applied to determine features showing a significant difference between factor levels. For features found to be significant, Tukey's Honest Significant Difference (HSD) was applied to determine between which levels the difference was significant ($p<0.05$). A False Discovery Rate (FDR) correction (Benjamini-Hochburg) was applied to adjust for multiple testing and control the number of false positives ($q < 0.05$) for both t-test and ANOVA.

***Software***

All peak matrix processing, univariate and multivariate analyses were performed in the R environment using STRUCT (STatistics in R Using Class Templates) and STRUCTToolbox packages, which make use of PMP and SBCMS packages. These packages are maintained by Phenome Centre Birmingham and available on GitHub (https://github.com/computational-metabolomics).

15. Dieterle F, Ross A, Schlotterbeck G, Senn H. Probabilistic quotient normalization as robust method to account for dilution of complex biological

mixtures. Application in 1H NMR metabonomics. Anal Chem **2006**;78(13):4281-90 doi 10.1021/ac051632c.

16. Parsons HM, Ludwig C, Gunther UL, Viant MR. Improved classification accuracy in 1- and 2-dimensional NMR metabolomics data using the variance stabilising generalised logarithm transformation. BMC Bioinformatics **2007**;8:234 doi 10.1186/1471-2105-8-234.

17. Sumner LW, Amberg A, Barrett D, Beale MH, Beger R, Daykin CA*, et al.* Proposed minimum reporting standards for chemical analysis Chemical Analysis Working Group (CAWG) Metabolomics Standards Initiative (MSI). Metabolomics **2007**;3(3):211-21 doi 10.1007/s11306-007-0082-2.

**Supplementary Table 1**

Gistic scores and mRNA expression levels for select genes in Tg(mitfa:BRAFV600E);p53 Zebrafish melanoma samples

**Supplementary Table 2**

GSEA analysis of significantly differentially expressed genes comparing either NRASG12D-positive EGFP-expressing (n=3) and NRASG12D-positive Dgat1a over-expressing (n=5) tumors or comparing DGAT1 mRNA high tumors (12/72) to the rest of the NRASmut TGCA dataset (60/72).

**Supplementary Table 3**

Positive ion total lipidomics data set for zebrafish tumours comparing either NRASG12D-positive EGFP-expressing (n=6) and NRASG12D-positive Dgat1a over-expressing (n=6) tumors.

**Supplementary Table 4**

Positive ion total lipidomics data set for DGAT1 overexpressing 888MEL melanoma cell line. 888MEL (n=3), 888MEL Clone 3 (n=3).

**Supplementary Table 5**

Negative ion total lipidomics data set for DGAT1 overexpressing 888MEL melanoma cell line. 888MEL (n=3), 888MEL Clone 3 (n=3).

**Supplementary Table 6**

Positive ion total lipidomics data set for DGAT1 inhibitor (A922500) treated A375 & SKMEL105 melanoma cell lines. A375 DMSO (n=3), A375 24h A922500 (n=3), A375 72h A922500 (n=3), SKMEL105 DMSO (n=3), SKMEL105 24h A922500 (n=3), SKMEL105 72h A922500 (n=3).

**Supplementary Table 7**

Negative ion total lipidomics data set for DGAT1 inhibitor (A922500) treated A375 & SKMEL105 melanoma cell lines. A375 DMSO (n=3), A375 24h A922500 (n=3), A375 72h A922500 (n=3), SKMEL105 DMSO (n=3), SKMEL105 24h A922500 (n=3), SKMEL105 72h A922500 (n=3).

**Supplementary Table 8**

Reagents and Resources

**Supplementary Figure 1 *DGAT1* amplification and up-regulation is associated with poor prognosis in melanoma**

(a) Cancer cell line encyclopaedia and TGCA data set accessed via cBioPortal quantifying alterations in indicated genes. (b) TCGA firehose legacy melanoma data set quantifying alterations in indicated genes via cBioPortal. (c) TGCA data set accessed via cBioPortal quantifying alterations in indicated genes. (d) Kaplan-Meier survival plots for co-amplified genes found on chromosome 8 in melanoma (top 25% *vs* bottom 75%, TCGA database). (e) G-Score of amplified regions of zebrafish chromosomes found in $BRAF^{V600E}$-positive; *tp53* mutant tumors, locations of human chromosome 8 zebrafish paralogues identified. (f) TGCA pan-cancer data set quantifying amplification of *DGAT1* accessed via cBioPortal.

**Supplementary Figure 2 Dgat1 functions as an oncoprotein in zebrafish**

(a) Hematoxylin and eosin stained transverse sections of EGFP expressing or Dgat1a over-expressing melanoma on the transgenic *mitfa*:$NRAS^{G12D}$; *tp53* mutant; *nacre* genetic background. (b) Principal component analysis (PCA) of RNA-seq data from $NRAS^{G12D}$-positive EGFP-expressing (n=3) and $NRAS^{G12D}$-positive Dgat1a over-expressing (n=5) tumors. (c) One minus pearson correlation of RNA-seq data NRASG12D-positive EGFP-expressing (n=3) and NRASG12D-positive Dgat1a over-expressing (n=5) tumors (morpheus). (d) RT-qPCR analysis of indicated genes in both $NRAS^{G12D}$-positive EGFP-expressing and Dgat1a over-expressing tumors. Relative expression calculated using a house keeping control gene (Mean). (e) TGCA pan cancer atlas RNA-seq data set filtered for $NRAS^{mut}$ melanoma patients.

Volcano plot where fold-change was calculated comparing *DGAT1* mRNA high tumors (12/72) to the rest of the NRAS$^{mut}$ data set (60/72). (f) Venn diagram of significantly altered gens comparing either NRASG12D-positive EGFP-expressing (n=3) and NRASG12D-positive Dgat1a over-expressing (n=5) tumors or comparing DGAT1 mRNA high tumors (12/72) to the rest of the NRASmut TGCA dataset (60/72).

**Supplementary Figure 3 DGAT1 activity is required for maintenance of S6K signalling**

(a) Pearson correlation between phospho-proteome samples following treatment of A375 cells with/without A922500 for 4 h (n=3). (b) Protein expression of phospho-S6 and phospho-eEF2 following treatment with/without DGAT1 inhibitor AZD3988. (c) Protein expression of phospho-S6 in indicated cell lines following treatment with/without DGAT2 inhibitor PF-06424439. (d) RT-qPCR analysis of DGAT2 gene expression in A375 cells following transfection with DGAT2 targeting siRNA (001, 002, 003, 004, pool) or scrambled control for 48 h (Mean ± SD, n=3) (upper). Corresponding protein expression of phospho-S6 (lower) (e) Protein expression of DGAT1, phospho-S6 and ERK (loading control) following transfection with DGAT1 overexpression vector or an empty vector control for 48 h. (f) Quantification of the percentage of 888MEL parental cells or Clone 3 DGAT1 over-expressing cells in S-phase following EdU incorporation (Mean ± SD, n>40). (g) Quantification of the population of cells in S-phase using EdU incorporation following transfection with DGAT1 targeting siRNA (007, 008, pool) or scrambled control for 48 h (Mean ± SD, n>5).

**Supplementary Figure 4 DGAT1 supports S6K dependent melanoma cell growth**

(a) Relative cell number determined by crystal violet staining following 72 h treatment with/without AZD3988 or A922500 (Mean ± SD, n>3). (b) Confluency curves determined by time-lapse microscopy in A375 cells following transfection with DGAT2 (001, 002, 003, 004, pool) or DGAT1 targeting scrambled siRNA using an Incucyte zoom system (Mean, n>3). (c) Quantification of the percentage of cells in S-phase using EdU incorporation following 24 h treatment with/without DGAT2 inhibitor (Mean ± SD, n>5). (d) Relative cell number of A375 cell line stained with crystal violet following over-expression of either a GFP control or constitutively active S6 kinase in the presence of either DGAT1 targeting siRNA (007, 008, pool) or a scrambled control for 48 h (Mean ± SD, n=3). (e) Protein expression of phospho-6 and phospho-P70S6K following treatment with/without DGAT1 inhibitor. Cells were grown in lipid restricted media for the duration of the experiment. (f) Relative cell number determined by crystal violet following transfection with DGAT1 overexpression vector or an empty vector control under normoxic or hypoxic conditions for 48 h. (1 % $O_2$) (upper) (Mean, n>3). Corresponding protein expression of DGAT1 (lower).

**Supplementary Figure 5 DGAT1-driven lipid droplets act as caretakers of mitochondrial health**

(a) UHPLC- lipidomic analysis of SKMEL105 cells following treatment with A922500 for 72 h and *NRAS*$^{G12D}$-positive Dgat1a-over-expressing (n=6) tumors showing the number of carbon-carbon double bonds in TAG species compared to the fold increase. Fold increase calculated as per figure 5A and figure 5D respectively. (b) Representative images and quantification of the number of lipid droplets per cell following BODIPY staining. A375 cells were treated with/without AZD3988 for 24 h (Mean ± SD, n>30). (c) Representative images and quantification of the number of lipid droplets per cell following BODIPY staining. Indicated cell lines were transfected with either DGAT1 targeting siRNA (007, 008, pool) or scrambled control for 48 h (Mean ± SD, n>30). (d) Representative images and quantification of the number of lipid droplets per cell following BODIPY staining. Indicated cells were transfected with either DGAT2 targeting siRNA (001, 002, 003, 004, pool) or a scrambled control for 48 h (Mean ± SD, n>30). (e) UHPLC- lipidomic analysis of A375 cells following treatment with/without A922500 for 24 or 72 h (showing lipid species that were annotated using MS/MS). Fold-change calculated relative to DMSO treated control. (f) A375 cells were stained with JC-1 dye following treatment with/without A922500 for 48 h and with/without Etomoxir for 4 h. The percentage of cells that lost red J-aggregates was calculated by using 1 μM CCP as a positive control for loss of mitochondrial membrane potential and comparing this to untreated cells to create two populations of cells in the flow cytometry analysis (Mean± SD, n>3). (g) Relative cell number as determined by crystal violet staining, indicted cell lines were treated with/without A922500 for 48 h and with/without Etomoxir for 24 h (Mean± SD, n=3).

**Supplementary Figure 6 DGAT1 suppression generates ROS leading to mitochondrial lipid peroxidation**

(a) Pearson correlation between total proteome samples following treatment of A375 cells with/without A922500 for 72 h (n=3). (b) Gene Ontology analysis of up-regulated genes (114) in A375 cells following treatment with A922500 for 72 h compared to DMSO treated control. Gene Ontology analysis was carried out using biological processes (enrichR) ranked by combined score and metascape ranked by log adjusted P value. (c) RT-qPCR analysis of indicated genes following treatment with/without A922500 for 24 or 72 h. Gene expression change calculated relative to DMSO control (Mean, n=3). (d) Quantification of ROS levels using dihydroethidium (DHE) fluorescence (ex 480nm, em 570nm) following treatment with/without A922500 for 48-72 h. Fold-change calculated relative to DMSO treated control (Mean, n>4). (e) Quantification of ROS levels using H2DCFDA (ex 480nm, em 535nm) in indicated cell lines following treatment with/without A922500 for 48-72 h. Fold-change calculated relative to DMSO treated control (Mean, n>4). (f) Indicated cell lines were stained with mitosox dye following treatment with A922500 for 48 h. Median fluorescence was determined using flow cytometry (Mean ± SD, n=6).

**Supplementary Figure 7 DGAT1 suppression triggers ROS induced apoptosis that is ameliorated by SESN2**

(a) Protein expression of SOD1 and SOD2 following treatment with/without A922500 for 24-72 h. (b) Cleaved caspase index in indicated cell lines following transfection with either a DGAT1 targeting siRNA (007, 008, pool) or a scrambled control determined by time-lapse microscopy using an Incucyte zoom system (Mean, n=3). (c) Cleaved caspase index in A375 cells following transfection with either a DGAT2 targeting (001, 002, 003, 004, pool), DGAT1 targeting or a scrambled siRNA determined by time-lapse microscopy using an Incucyte zoom system (Mean, n=3) (left). Corresponding protein expression of cleaved caspase3 following transfection with either a DGAT2 targeting siRNA (001, 002, 003, 004, pool), DGAT1 targeting siRNA (007, 008, pool) or a scrambled control (right). (d) Quantification of ROS levels using dihydroethidium fluorescence (ex 480nm, em 570nm) following treatment with/without a DGAT2 inhibitor for 24-72 h. Fold-change calculated relative to DMSO treated control (Mean, n>3). (e) Protein expression of Sestrin 2 and cleaved caspase3 following transfection with SESN2 targeting siRNA or a scrambled control and treatment with/without A922500 for 24 h. (f) Relative SESN2 expression in A375 cells transfected with SESN2 targeting siRNA (19,20) or a scrambled control followed by treatment with A922500 for 24 h. Fold-change calculated relative to DMSO treated control (Mean ± SD, n=3). (g) Protein expression of Sestrin 2 and phospho-S6 in indicated cell lines following transfection with SESN2 targeting siRNA or a scrambled control and treatment with/without A922500 for 2 h. (h) Model depicting the role of DGAT1 in melanoma. Figure made in Biorender.

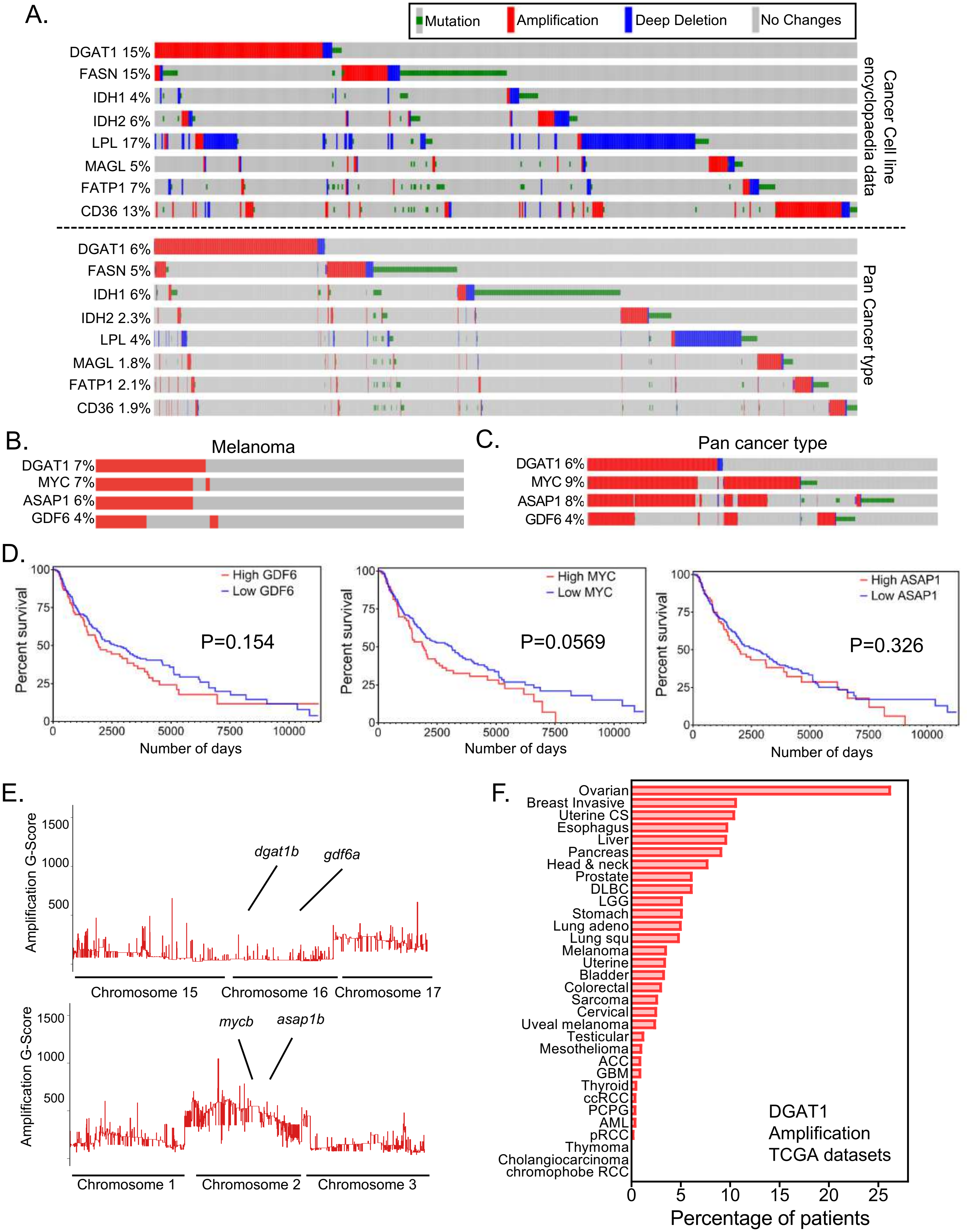


**Supplementary Figure 1** ***DGAT1*** **amplification and up-regulation is associated with poor prognosis in melanoma**

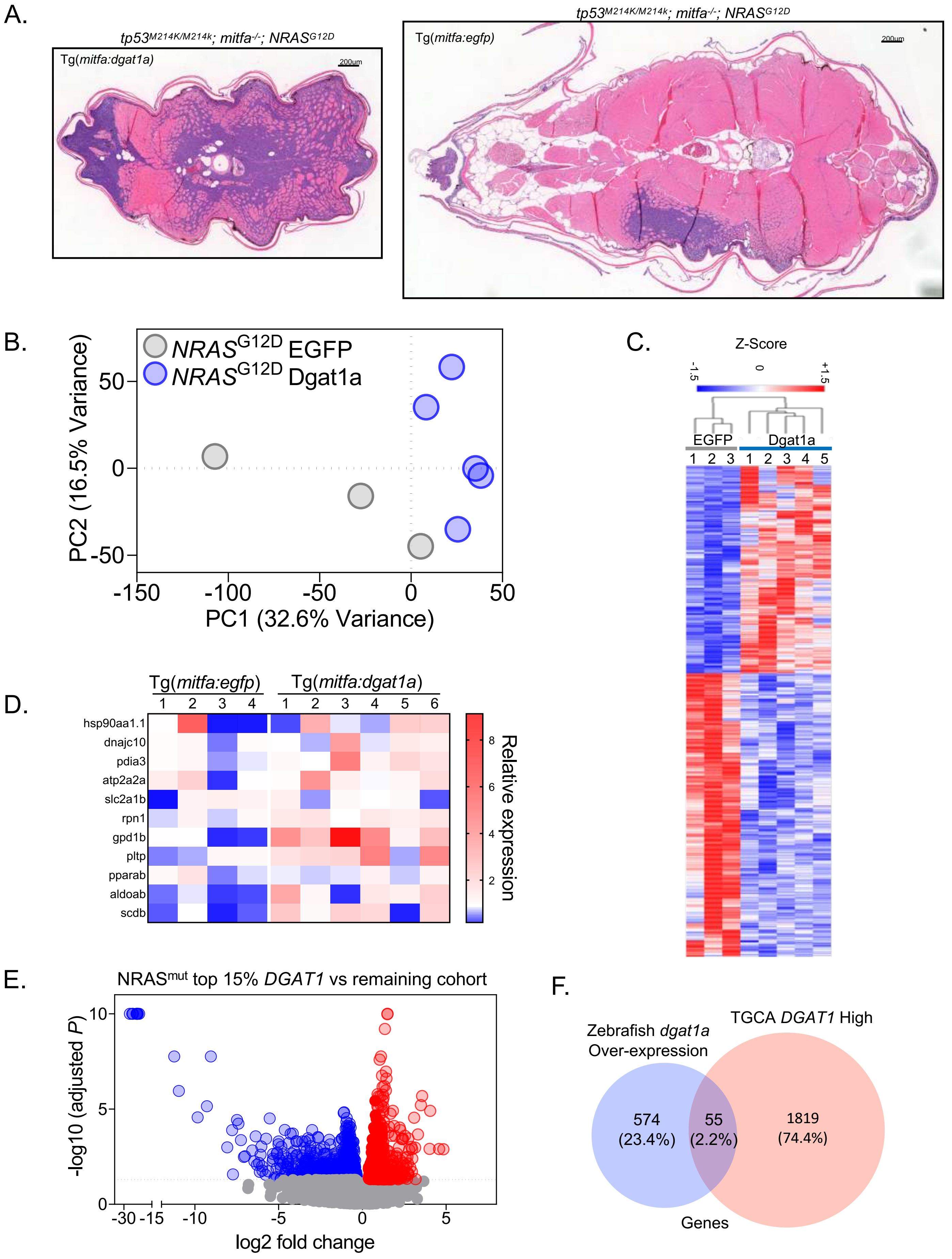


**Supplementary Figure 2 Dgat1 functions as an oncoprotein in zebrafish**

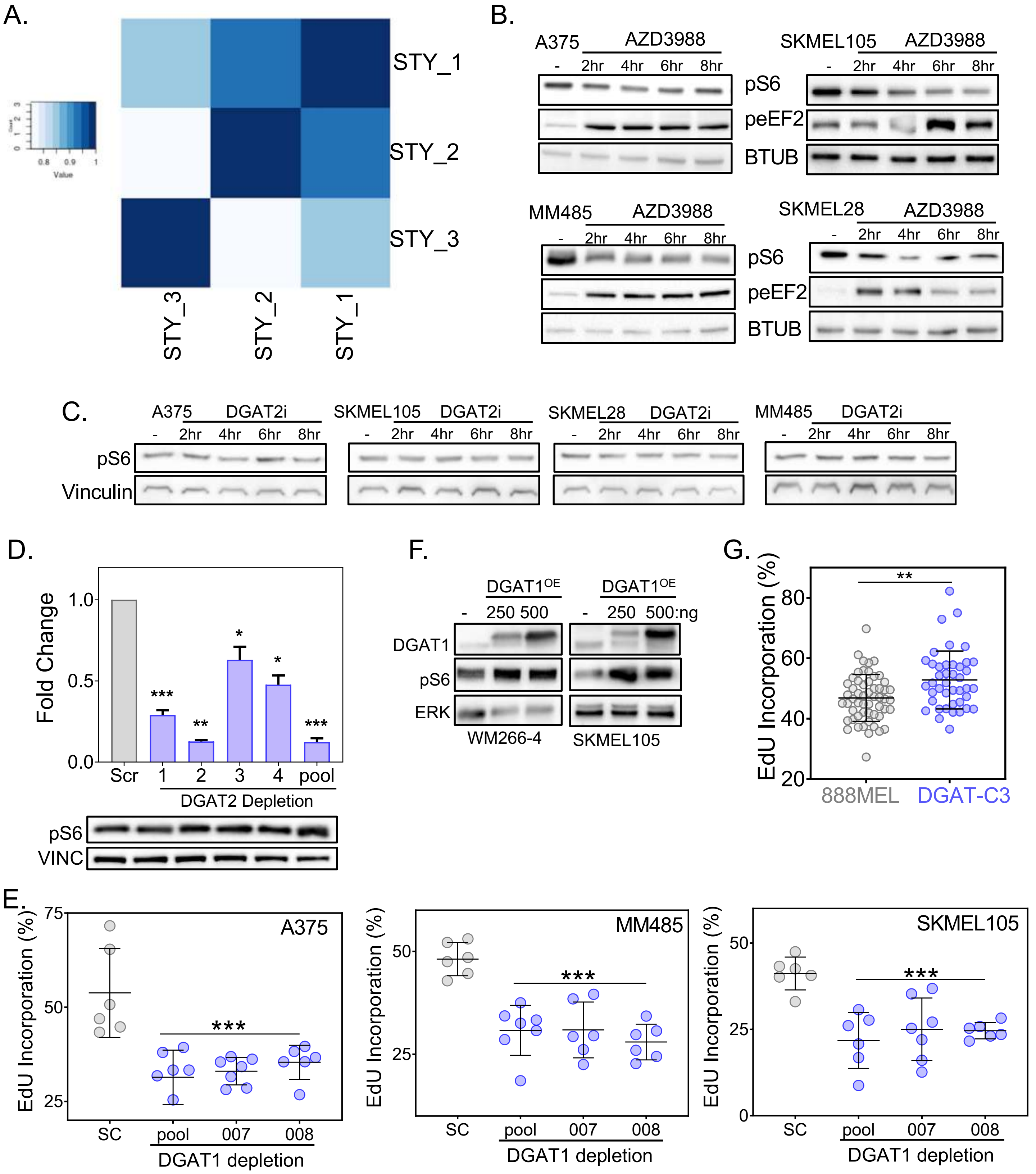


**Supplementary Figure 3 DGAT1 activity is required for maintenance of S6K signalling**

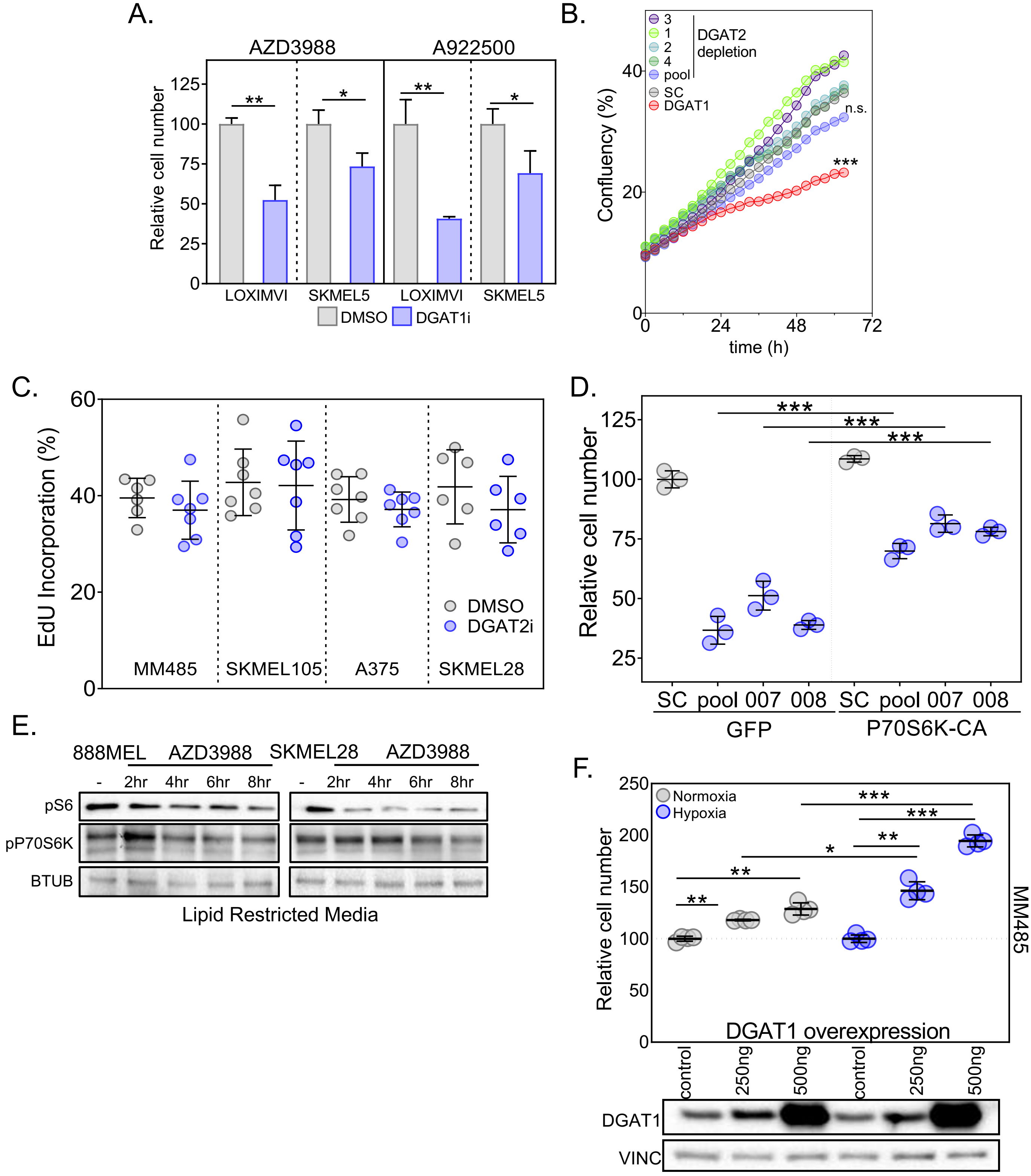


**Supplementary Figure 4 DGAT1 supports S6K dependent melanoma cell growth**

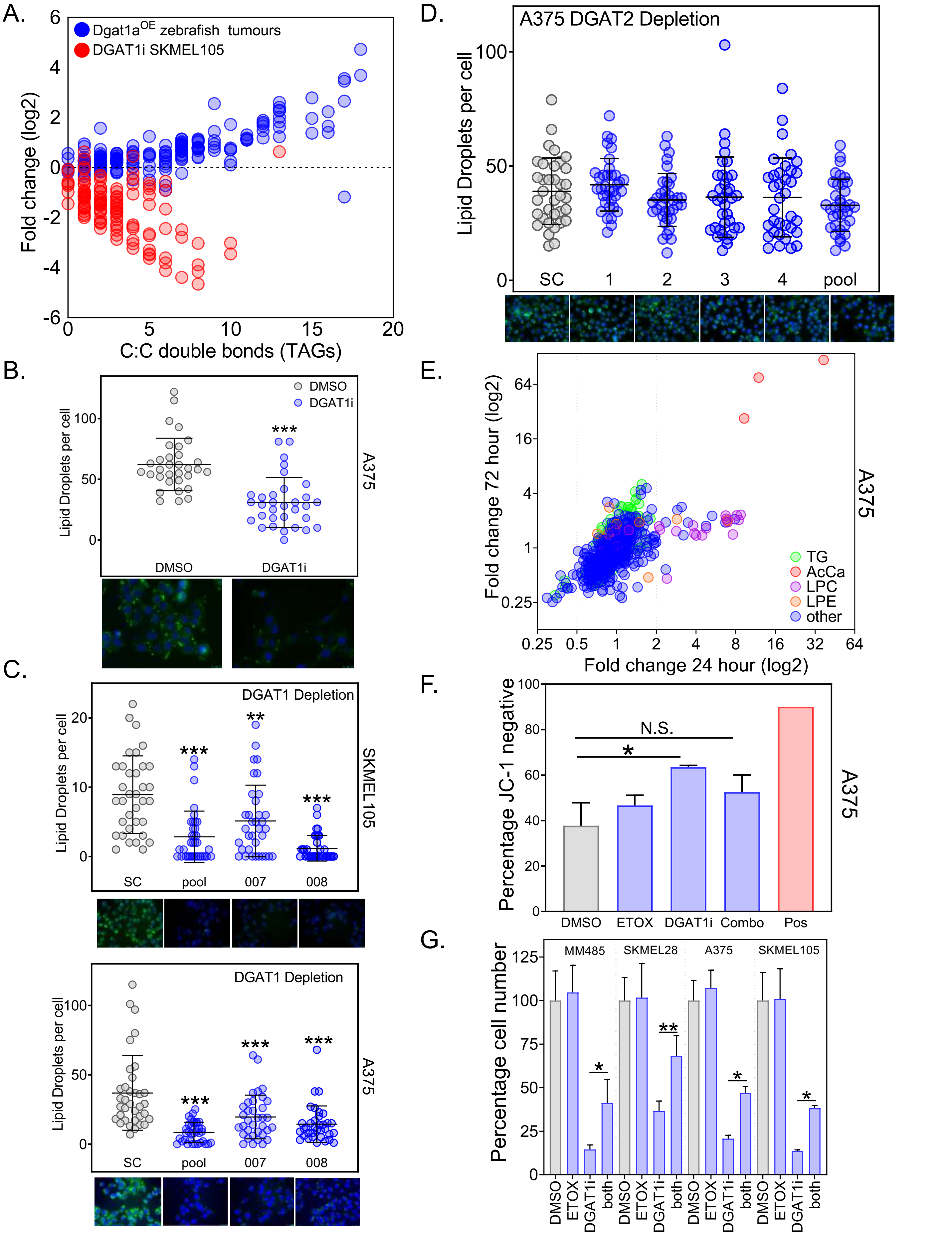


**Supplementary Figure 5 DGAT1-driven lipid droplets act as caretakers of mitochondrial health**

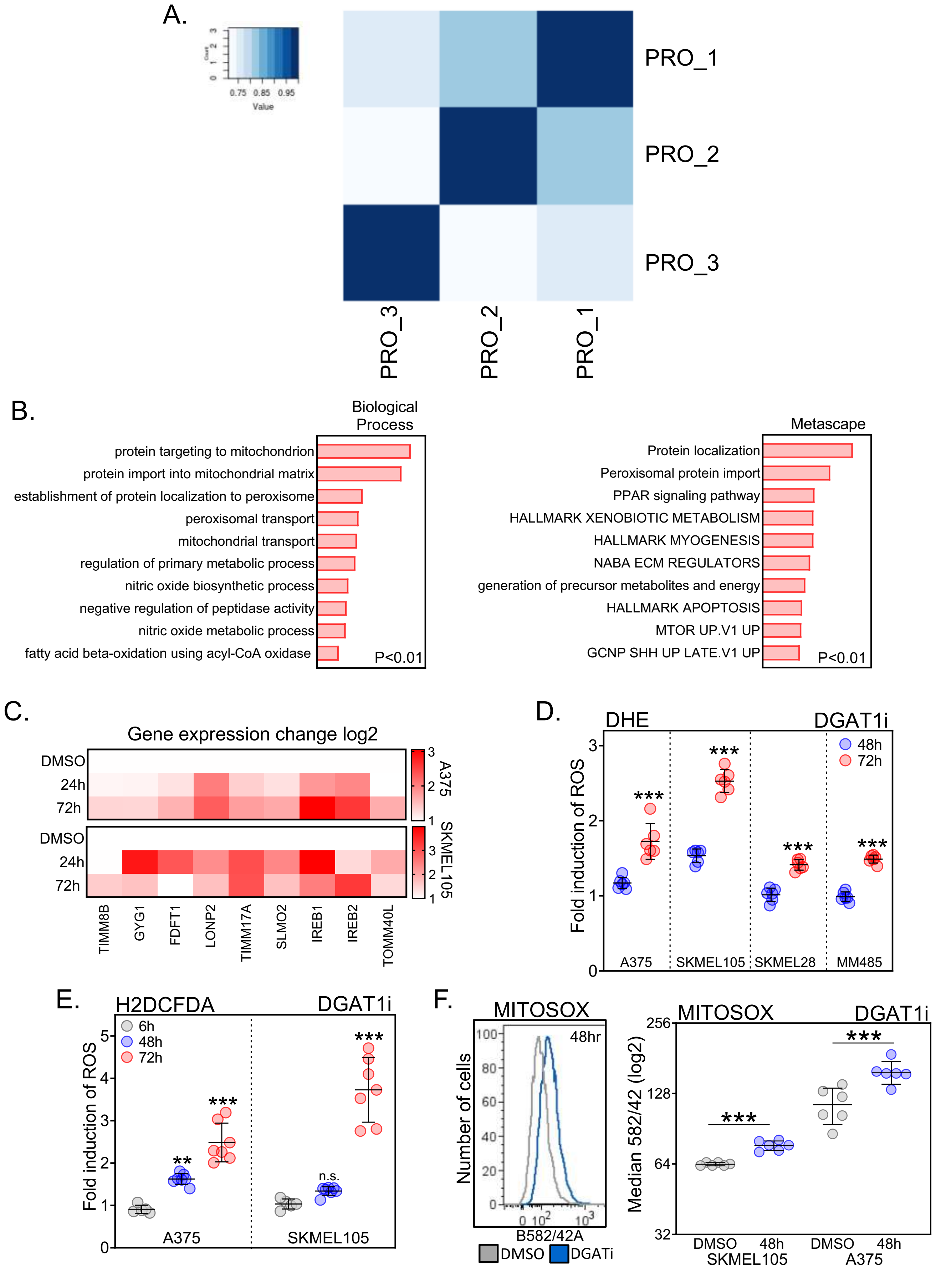


**Supplementary Figure 6 DGAT1 suppression generates ROS leading to mitochondrial lipid peroxidation**

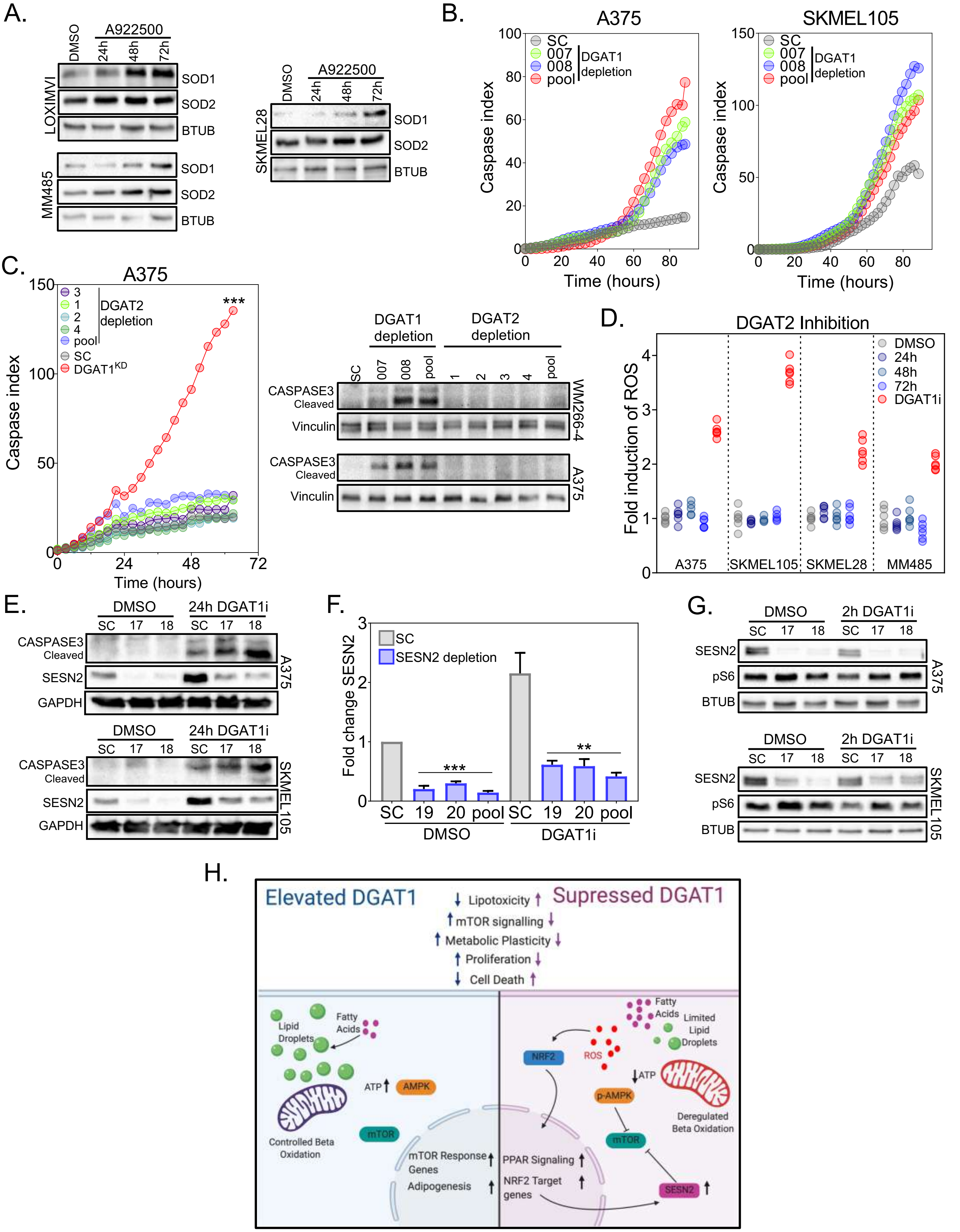


**Supplementary Figure 7 DGAT1 suppression triggers ROS induced apoptosis that is ameliorated by SESN2**
